# An image-computable model on how endogenous and exogenous attention differentially alter visual perception

**DOI:** 10.1101/2021.01.26.428173

**Authors:** Michael Jigo, David J. Heeger, Marisa Carrasco

**Author notes:** Corresponding author information: Michael Jigo.

## Abstract

Attention alters perception across the visual field. Typically, endogenous (voluntary) and exogenous (involuntary) attention similarly improve performance in many visual tasks, but they have differential effects in some tasks. Extant models of visual attention assume that the effects of these two types of attention are identical and consequently do not explain differences between them. Here, we develop a model of spatial resolution and attention that distinguishes between endogenous and exogenous attention. We focus on texture-based segmentation as a model system because it has revealed a clear dissociation between both attention types. For a texture for which performance peaks at parafoveal locations, endogenous attention improves performance across eccentricity, whereas exogenous attention improves performance where the resolution is low (peripheral locations) but impairs it where the resolution is high (foveal locations) for the scale of the texture. Our model emulates sensory encoding to segment figures from their background and predict behavioral performance. To explain attentional effects, endogenous and exogenous attention require separate operating regimes across visual detail (spatial frequency). Our model reproduces behavioral performance across several experiments and simultaneously resolves three unexplained phenomena: (1) the parafoveal advantage in segmentation, (2) the uniform improvements across eccentricity by endogenous attention and (3) the peripheral improvements and foveal impairments by exogenous attention. Overall, we unveil a computational dissociation between each attention type and provide a generalizable framework for predicting their effects on perception across the visual field.

## INTRODUCTION

Endogenous and exogenous spatial attention prioritize subsets of visual information and facilitate their processing without concurrent eye movements (1–3). Selection by endogenous attention is goal-driven and adapts to task demands whereas exogenous attention transiently and automatically orients to salient stimuli (1–3). In most visual tasks both types of attention typically improve visual perception similarly (e.g., acuity (4–6), visual search (7, 8), perceived contrast (9–11)). Consequently, models of visual attention do not distinguish between endogenous and exogenous attention (e.g., (12–19)). However, stark differences also exist. Each attention type differentially modulates neural responses (20, 21) and fundamental properties of visual processing, including temporal resolution (22, 23), texture sensitivity (24), sensory tuning (25), contrast sensitivity (26) and spatial resolution (27–34).

The effects of endogenous and exogenous attention are dissociable during texture segmentation, a visual task constrained by spatial resolution (reviews(1–3)). Whereas endogenous attention optimizes spatial resolution to improve the detection of an attended texture (32–34), exogenous attention reflexively enhances resolution even when detrimental to perception (27–31, 34). Extant models of attention do not explain these well-established effects.

Two main hypotheses have been proposed to explain how attention alters spatial resolution. Psychophysical studies ascribe attentional effects to modulations of spatial frequency (SF) sensitivity (30, 33). Neurophysiological (13, 35, 36) and neuroimaging (37, 38) studies bolster the idea that attention modifies spatial profiles of neural receptive fields (2). Both hypotheses provide qualitative predictions of attentional effects but do not specify their underlying neural computations.

Differences between endogenous and exogenous attention are well established in segmentation tasks and thus provide an ideal model system to uncover their separate roles in altering perception. Texture-based segmentation is a fundamental process of mid-level vision that isolates regions of local structure to extract figures from their background (39–41). Successful segmentation hinges on the overlap between the visual system’s spatial resolution and the levels of detail (i.e., SF) encompassed by the texture (39, 41, 42). Consequently, the ability to distinguish between adjacent textures varies as resolution declines toward the periphery (43–46). Each attention type differentially alters texture segmentation, demonstrating that their effects shape spatial resolution (reviews(1–3)).

Current models of texture segmentation do not explain performance across eccentricity and the distinct modulations by attention. Conventional models treat segmentation as a feedforward process that encodes the elementary features of an image (e.g., SF and orientation), transforms them to reflect the local structure (e.g., regions of similarly oriented bars), then pools across space to emphasize texture-defined contours (39, 41, 47). Few of these models account for variations in resolution across eccentricity (46, 48, 49) or endogenous (but not exogenous) attentional modulations (18, 50). All others postulate that segmentation is a ‘preattentive’ (42) operation whose underlying neural processing is impervious to attention (39, 41, 46–49).

Here, we develop a computational model in which feedforward processing and attentional gain contribute to segmentation performance. We augment a conventional model of texture processing (39, 41, 47). Our model varies with eccentricity and includes contextual modulation within local regions in the stimulus via normalization (51), a canonical neural computation (52). The defining characteristic of normalization is that an individual neuron is (divisively) suppressed by the summed activity of neighboring neurons responsive to different aspects of a stimulus. We model attention as multiplicative gains (attentional gain factors (15)) that vary with eccentricity and SF. Attention shifts sensitivity toward fine or coarse spatial scales depending on the range of SFs enhanced.

Our model is image-computable, which allowed us to reproduce behavior directly from grayscale images used in psychophysical experiments (6, 26, 27, 29–33). The model explains three signatures of texture segmentation hitherto unexplained within a single computational framework (Figure 1). (i) The central performance drop (CPD) (27-34, 43-46) (Figure 1A), i.e., the parafoveal advantage of segmentation over the fovea. (ii) The improvements in the periphery and impairments at foveal locations induced by exogenous attention (27–32, 34) (Figure 1B). (iii) The equivalent improvements across eccentricity by endogenous attention (32–34) (Figure 1C).

**Figure 1.**
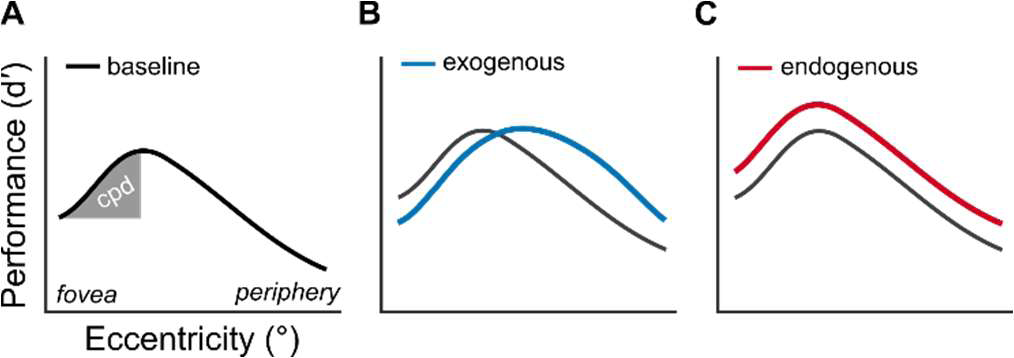
Signatures of texture segmentation. (A) Central performance drop. Shaded region depicts the magnitude of the central performance drop. Identical axis labels are omitted in panels B and C. (B) Exogenous attention modulation. Exogenous attention improves segmentation performance in the periphery and impairs it near the fovea. (C) Endogenous attention modulation. Endogenous attention improves segmentation performance across eccentricity.

Whereas our analyses focused on texture segmentation, our model is general and can be applied to other visual phenomena. We show that the model predicts contrast sensitivity across SF and eccentricity as well as the effects of attention on contrast sensitivity and acuity; i.e. in tasks in which both endogenous and exogenous attention have similar or differential effects on performance. To preview our results, model comparisons revealed that normalization is necessary to elicit the CPD and that separate profiles of gain enhancement across SF (26) generate the effects of exogenous and endogenous attention on texture segmentation. A preferential high-SF enhancement reproduces the impairments by exogenous attention due to a shift in visual sensitivity toward details too fine to distinguish the target at foveal locations. The transition from impairments to improvements in the periphery results from exogenous attentional gain gradually shifting to lower SFs that are more amenable for target detection. Improvements by endogenous attention result from a uniform enhancement of SFs that encompass the target, optimizing visual sensitivity for the attended stimulus across eccentricity.

## RESULTS

### Image-computable model of attention and spatial resolution

We developed an observer model based on established principles of neural computation (51, 52), pattern (53, 54) and texture vision (39, 41, 47) and attentional modulation (15). The model incorporates elements of the Reynolds-Heeger normalization model of attention (NMA) (15) and illuminates how attention alters contrast and texture sensitivity across SF and eccentricity. We implement: (i) SF-tuned gain modulation to emulate the decline in contrast sensitivity and peak SF preference with eccentricity. (ii) Spatial summation of normalized inputs to generate texture selectivity. (iii) Separate attentional gain profiles across SF to reproduce effects of exogenous and endogenous attention. The model is composed of four components: stimulus drive, attentional gain, suppressive drive and spatial summation (Figure 2A). Following NMA, attention adjusts the gain on the stimulus drive before normalization. For a full description of the model, see Methods.

**Figure 2.**
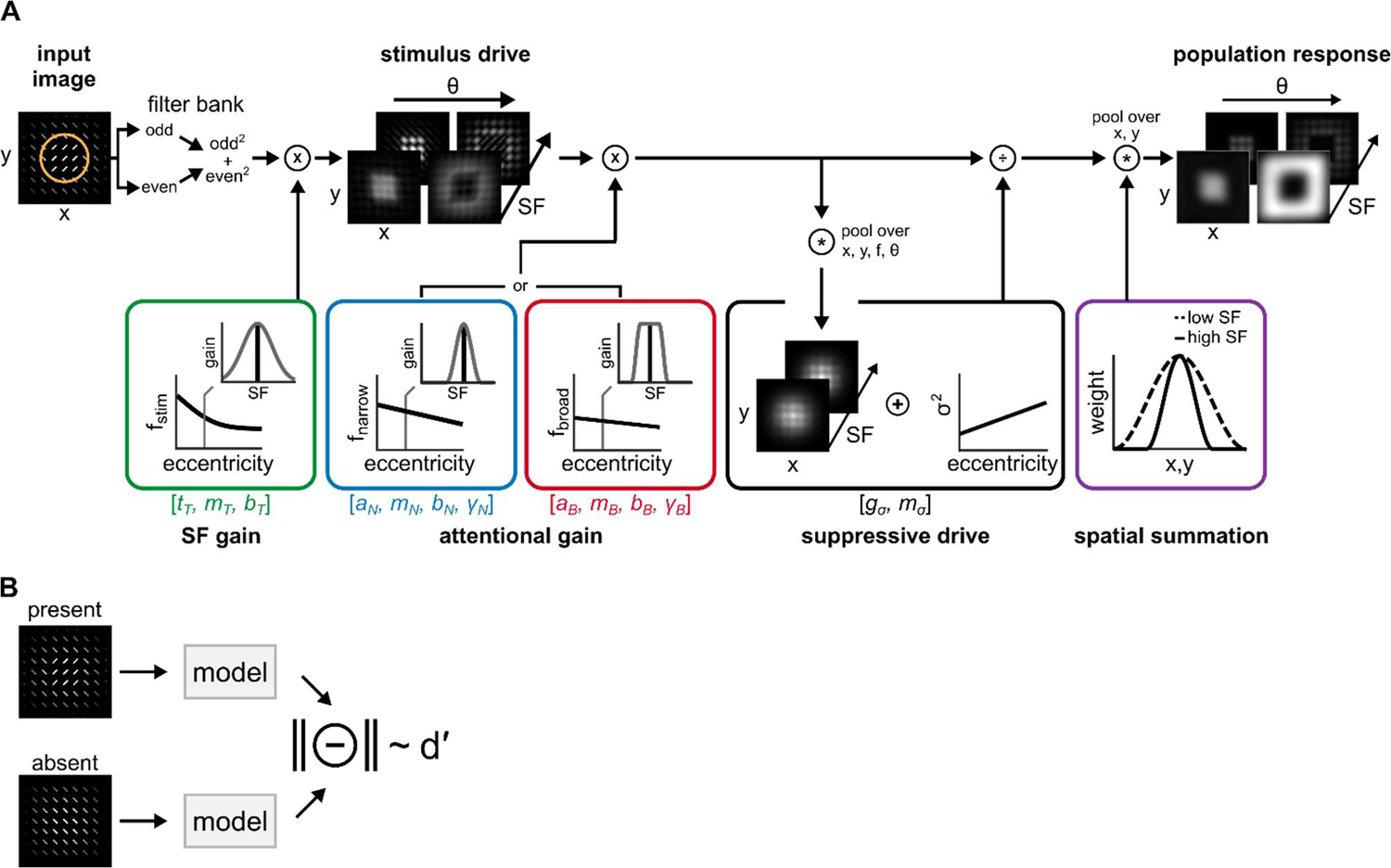
Image computable model of attention and spatial resolution (A) Model structure. A filter bank of linear receptive fields decomposes an image. Filter responses are squared and summed across quadrature-phase pairs (odd, even), yielding contrast energy outputs. SF gain scales contrast energy across SF and eccentricity (green box). The solid black line depicts the center frequency of the tuning function (f_stim_); insets display the full SF tuning function at a single eccentricity. The stimulus drive characterizes contrast energy at each pixel in the image, filtered through feature-selective and eccentricity-dependent receptive fields. Attentional gain multiplicatively scales the stimulus drive at a circumscribed region within the image (orange circle in left panel) and varies across SF and eccentricity. The center SF of attentional gain varies with eccentricity (solid black lines in blue and red boxes). Across SF, attentional gain follows either a narrow profile (blue box) or a broad profile (red box), each centered on a given frequency (f_narrow_ or f_broad_). The suppressive drive comprises the attention-scaled stimulus drive pooled across a local neighborhood of positions, SFs and uniformly across orientation. Contrast gain, σ^2^, adjusts suppression magnitude across eccentricity. Spatial summation follows normalization (purple box) and generates the population response. Pooling area varies inversely with SF tuning. Variables displayed within the square brackets depict model parameters fit to behavior. (B) Target discriminability. Population responses for texture images with (present) or without (absent) a target patch are computed. The vector magnitude of their difference produces a metric proportional to d′, assuming independent and identically distributed Gaussian output noise.

#### Stimulus drive

We simulate bottom-up responses of a collection of linear receptive fields (RFs), each jointly tuned to spatial position, SF and orientation. Images are processed through a filter bank (55) covering the visual field at several SFs and orientations using bandwidths compatible with neurophysiological (54) and psychophysical (53) measurements. Filter outputs are combined across quadrature phase (56), yielding contrast energy images corresponding to different SFs and orientations. These outputs simulate the responses of complex cells in primary visual cortex (54, 56). The gain on individual RFs varies as a function of SF and eccentricity preference (Figure 2A, green). Following the behavior of individual neurons (54) and pattern vision (53), gain modulation is narrowly tuned to high SFs near the fovea and progressively shifts to low SFs with eccentricity. Consequently, the stimulus drive reflects local spectral energy within each patch in an image, filtered through feature-selective RFs that vary with eccentricity.

#### Attentional gain

Attention is implemented as a gain control mechanism that scales the gain on the stimulus drive (15). The magnitude of attentional gain is largest at the cued location (Figure 2A, orange) and varies with the eccentricity and SF preference of each RF. Motivated by findings of psychophysical experiments that manipulated endogenous and exogenous attention (26), two SF-tuned profiles are assessed—narrow and broad. The narrow profile selectively enhances a small range of SFs at each eccentricity (Figure 2A, blue); the broad profile uniformly enhances SFs (Figure 2A, red).

#### Suppressive drive

Suppression operates via divisive normalization (51, 52). Normalized responses are proportional to the attention-scaled stimulus drive divided by a normalization pool plus a constant σ^2^ that increases with eccentricity. This constant adjusts the model’s overall sensitivity to contrast (i.e., contrast gain; Figure 2A, black). The normalization pool consists of the attention-scaled stimulus drive across nearby spatial locations (surround suppression (57)), uniformly across orientation (cross-orientation suppression (58)) and across preferred and neighboring SFs (cross-frequency suppression (59)) of individual RFs. Such broad suppressive pools are supported by physiological (57, 58, 60) and psychophysical (59, 61, 62) findings and models of visual processing (51).

#### Spatial summation

Normalized responses are weighted and summed across space within each SF and orientation filter. Spatial summation followed normalization (63), which accentuated texture-defined contours within the image. The size of pooling regions scale with the SF preference of each RF (39, 41) (Figure 2A, purple); larger for low than for high SFs. This implements an inverse relation between the integration area of individual RFs and their SF tuning.

#### Target discriminability

The model generated measures of discriminability (d′) in a texture segmentation task (Figure 2B). The model generated population responses to two texture images. One contained a target patch whose orientation differed from its surround (target-present) and the other consisted of uniform orientation throughout (target-absent). The vector length (i.e., Euclidean norm) of the difference between population responses indexed d′. This measure is proportional to behavioral performance, assuming the addition of normally distributed noise after normalization.

### Texture stimuli, behavioral protocol and optimization strategy

Stimuli. Model parameters were constrained by data from ten published psychophysical experiments. Exogenous attention was manipulated in six (27, 29–32) (Figure 3A-F) and endogenous attention in four experiments (32, 33) (Figure 3G-J). In each experiment, observers distinguished a patch of one orientation embedded within a background of differing orientation at several possible eccentricities.

**Figure 3.**
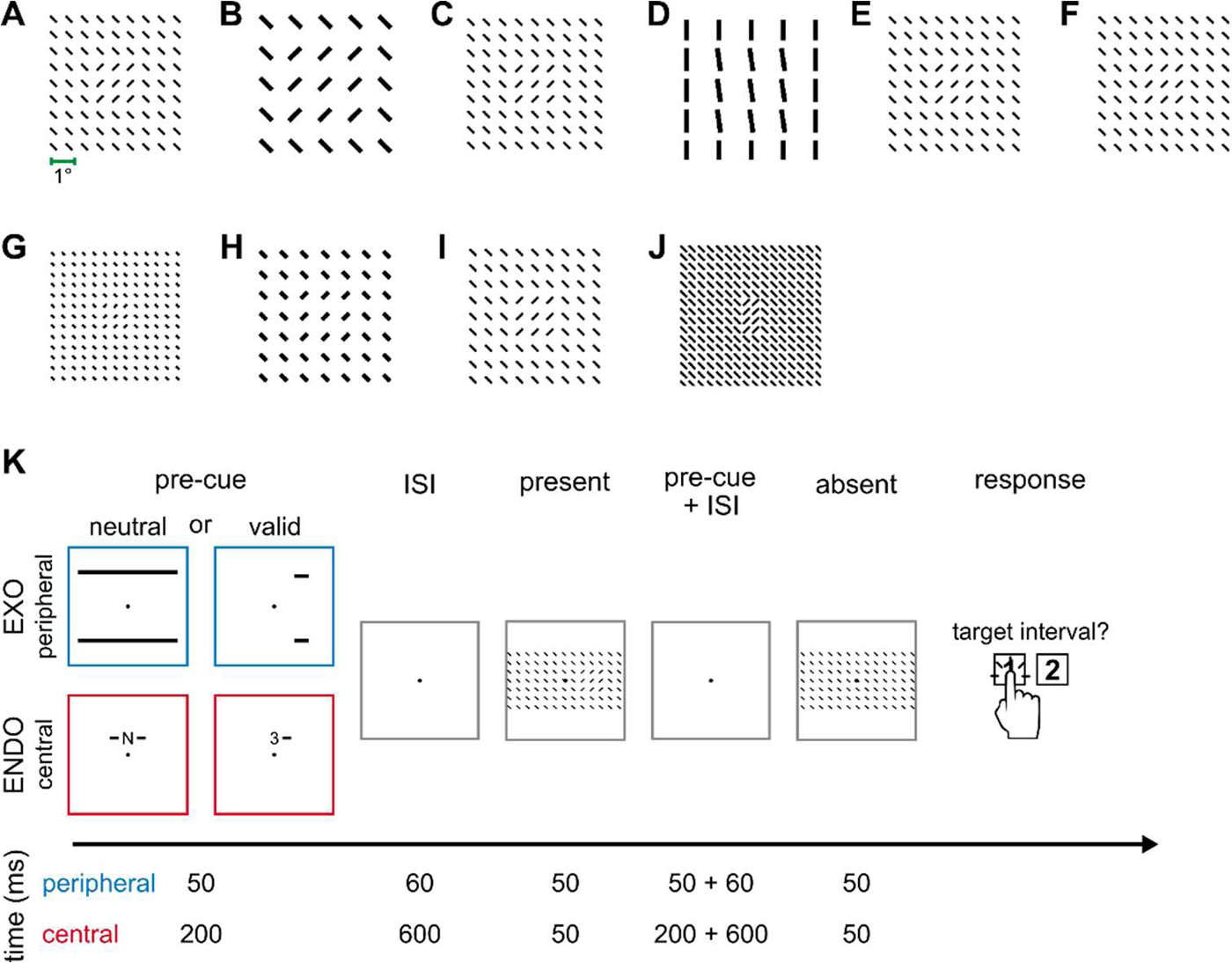
Texture stimuli and a typical texture segmentation behavioral protocol. Target-present texture stimuli used in (A-F) exogenous attention and (G-J) endogenous attention experiments, displayed at their respective spatial scales. Textures displayed include: (A) Fine and (B) coarse-scale textures used in Yeshurun & Carrasco, 1998 (27); (C) Talgar & Carrasco, 2002 (29) with targets placed on the vertical meridian; (D) Carrasco, Loula & Ho, 2006 (30) wherein observers discriminated the target’s orientation; (E) Yeshurun & Carrasco, 2008 (31) where the cue’s size was manipulated; (F) Experiment 2 of Yeshurun, Montagna & Carrasco, 2008 (32) with targets placed on the horizontal meridian; (G) Experiment 1 of Yeshurun, Montagna & Carrasco, 2008 (32) with targets placed on the horizontal meridian; (H) Experiment 3 and (I) Experiment 4 of Yeshurun, Montagna & Carrasco, 2008 (32) wherein fine and coarse-scale textures were displayed, respectively; and (J) Barbot & Carrasco, 2017 (33) with targets placed on the intercardinal meridians. (K) Two-interval forced choice protocol typically used to assess texture segmentation performance. EXO corresponds to exogenous attention and ENDO to endogenous attention. Numbers denote the representative timing information for each pre-cue—peripheral (blue) and central (red)—and their corresponding inter-stimulus intervals (ISI). Neutral pre-cues equally distributed attention to all possible target locations. Valid peripheral pre-cues appeared near the upcoming target location whereas valid central pre-cues symbolically indicated the upcoming target location. In the displayed example, the number “3” and the adjacent line indicate that the target would appear at a peripheral eccentricity in the right visual hemifield.

#### Behavioral protocol

Performance was typically measured with a two-interval forced choice protocol (Figure 3K). Observers maintained fixation at the display’s center while viewing two intervals of texture stimuli, one of which randomly contained a target texture. Different pre-cues at their optimal timing manipulated exogenous or endogenous attention. Brief peripheral pre-cues manipulated exogenous attention and appeared before both intervals, but near the upcoming target location in the interval containing the target (27–32, 34). Symbolic pre-cues manipulated endogenous attention. Pre-cues appeared near fixation and indicated the target location in the target-present interval (32, 33). Attention effects were determined relative to a neutral condition, in which observers distributed attention across all possible target locations. Behavioral performance displayed the three signatures of texture segmentation: (i) The CPD emerged in the neutral condition (Figure 1A). (ii) Peripheral pre-cues improved performance in the periphery and impaired it at foveal locations (Figure 1B). (iii) Central, symbolic pre-cues improved performance at all eccentricities (Figure 1C).

#### Optimization

To identify the computations that underlie each signature, we separately fit the model to three subsets of behavioral data. First, the CPD was isolated from attentional effects by fitting to the neutral condition from all ten experiments. Second, exogenous attentional effects were assessed by fitting to neutral and peripheral cueing conditions from the six exogenous attention experiments. Third, endogenous attentional effects were assessed by fitting to neutral and central cueing conditions from the four endogenous attention experiments. The model was jointly fit to each subset of data, with model parameters shared among experiments within a subset (Table S2-S4).

### Contextual modulation and spatial summation mediate the CPD

To identify the computations mediating the CPD, we fit the model to group-average performance across all experiments’ neutral condition (103 data points). 15 model parameters constrained performance (Table S2). To account for differences in contrast sensitivity due to variable display properties among experiments (e.g., mean luminance), foveal contrast gain (gσ; Figure 2A) was independently determined for each of ten experiments (10 parameters). Two separate parameters determined foveal SF preference (t_T_)–one shared among exogenous attention studies and another among endogenous attention studies. The remaining three parameters–SF bandwidth (b_T_), the gradual increase in contrast gain (mσ) and the progressive shift to lower SFs with eccentricity (m_T_)– were shared among all experiments. Attentional gain was not included for these fits.

The model reproduced the CPD and its dependence on texture scale (Figure 4). For a fine-scale texture—characterized by narrow, densely spaced lines—performance peaked within the parafovea (4 deg) and declined toward the fovea and periphery (Figure 4A). Differences between target-present and target-absent stimuli were largest within the 2 cpd filter (Figure 4A, middle).

**Figure 4.**
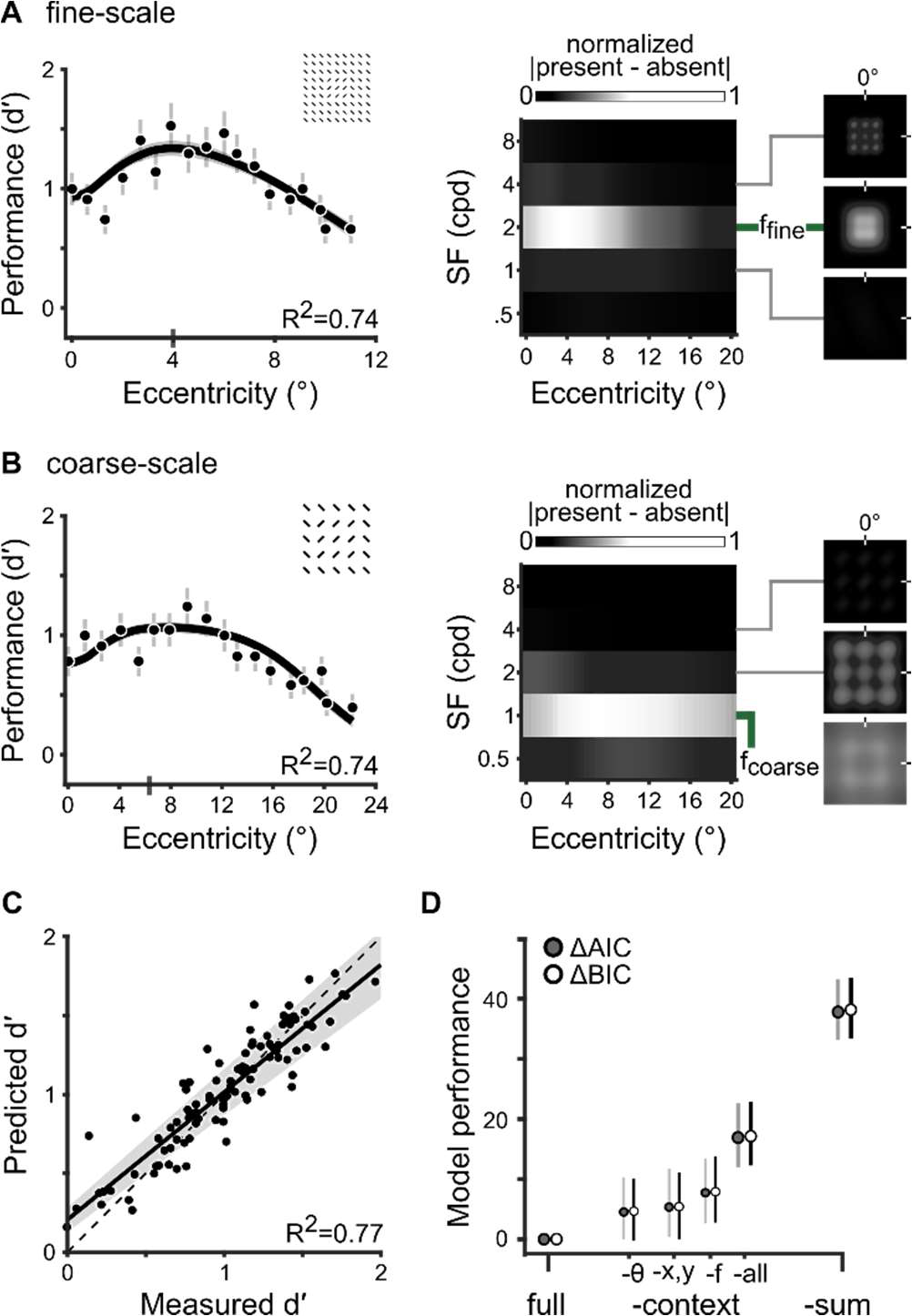
Contextual modulation and spatial summation mediate the CPD (A) Left. Fit to Experiment 1 in Yeshurun & Carrasco, 1998 (27). Dots (n=18) and error bars depict group-average performance and ±1 SEM. The black line and shaded regions depict the median and 68% bootstrapped confidence interval of model fits. The gray vertical bar on the x-axis indicates the eccentricity of peak performance. The inset shows the textures stimulus. Middle. The matrix depicts the absolute value of differences between target-present and target-absent population responses, normalized by the maximum across eccentricity and averaged across orientation and space. f_fine_ denotes the SF filter with the largest difference between population responses. We use absolute differences only to visualize the SFs that drove discriminability. Right. Spatial distribution of the absolute value of differences between target-present and target-absent population responses. Each panel depicts a subset of receptive fields centered on the fovea and tuned to one of three SFs (4, 2, 1 cpd) and an orientation of 30°. (A) (B) Fit to Experiment 2 (n=18) in Yeshurun & Carrasco, 1998 (27). The model jointly fits neutral performance with parameters shared among all ten experiments, including the data shown in A. Visualization follows the conventions in A. Note that eccentricity (x-axis) is twice that of A. f_coarse_ denotes the SF filter that best distinguished the coarse-scale target. (B) Goodness-of-fit for the neutral condition across ten experiments (n=103). Each dot depicts the measured (x-axis) and predicted (y-axis) performance at a given eccentricity. The solid line and shaded area depict the best-fitting regression line and its 95% confidence interval. The dashed line indicates the unity line y=x. (C) Model comparisons using AIC and BIC. Positive values indicate models underperforming, relative to the full model. ‘-context’ describes restrictions of contextual modulation: ‘-θ’ denotes the variant without cross-orientation suppression, ‘-f’ without cross-frequency suppression, ‘-x,y’ without surround suppression and‘-all’ devoid of all contextual modulation. ‘-sum’, denotes the model variant without spatial summation. The dots and error bars denote the median and 95% confidence interval of the bootstrap distribution.

This filter best differentiated the target patch from a homogenous texture; we denote its center SF as f_fine_. A coarser texture was best distinguished by lower SFs (1 cpd, f_coarse_), which exaggerated the CPD, moving peak performance to a farther eccentricity (∼6 deg; Figure 4B). The CPD was well-fit in all experiments (Figure 4C); 77% of the variance was explained (95% bootstrapped CI = [70 80]), with the best-fitting regression line falling close to the unity line.

Previous models qualitatively matched the CPD through spatial summation (46, 48, 49), but ignored the contributions of contextual modulation via normalization. To assess the contribution of each operation to behavior, we compared the full model to variants that either lacked components of the suppressive drive (cross-orientation, cross-frequency, and/or surround suppression) or spatial summation (Figure 4D). We restricted contextual modulation (-context) by separately limiting the pool of orientations (-θ), SFs (-f), spatial positions (-x,y) or all simultaneously (-all) such that suppressive modulations due to featural attributes and/or spatial positions outside each receptive field’s tuning were removed. The final variant lacked spatial summation (-sum), which resulted in a population response that consisted of only normalized inputs. Removing spatial summation attenuates the response to regions of similar orientation (e.g., target patch). Each model was fit to behavioral performance in the neutral condition across all experiments and compared using Akaike Information Criterion (AIC) (64) and Bayesian Information Criterion (BIC) (65).

Removing contextual modulation or spatial summation attenuated the CPD (Figure S1). We measured model performance relative to the full model, which yielded ΔAIC and ΔBIC scores; positive values represent a decrease in model performance. We use “M” and “CI” to denote the median and 95% confidence interval of the bootstrapped distribution. Model performance fell without cross-orientation suppression (ΔAIC: M=4.8, CI=[-0.1 9.7]; ΔBIC: M=4.6, CI=[-0.2 9.6]), cross-frequency suppression (ΔAIC: M=7.9, CI=[2.7 13.2]; ΔBIC: M=7.7, CI=[2.4 13.8]), surround suppression (ΔAIC: M=5.4, CI=[0.03 11.0]; ΔBIC: M=5.4, CI=[-0.1 11.5]), and without all forms of contextual modulation (ΔAIC: M=17.0, CI=[11.5 22.1]; ΔBIC: M=16.9, CI=[11.6 22.4]). Without spatial summation, model performance decreased as well (ΔAIC: M=37.8, CI=[33.3 42.6]; ΔBIC: M=37.8, CI=[33.1 42.8]). Thus, reliable reproduction of the CPD requires both contextual modulation and spatial summation.

### Narrow high-SF enhancement generates exogenous attention effects

The model predicted behavior in neutral and peripheral cueing conditions across six experiments (146 data points). Exogenous attention was modeled as a narrow SF gain profile (Figure 2, blue), motivated by psychophysical measurements (26). 14 free parameters constrained model behavior (Table S3). Model parameters that determined neutral cueing performance—foveal contrast gain (gσ), SF tuning (t_T_), SF bandwidth (b_T_), the increase in contrast gain (mσ) and the decline in SF preference with eccentricity (m_T_)—were configured identically as described above. Four parameters, shared among experiments, determined attentional gain–foveal SF preference (a_N_), the gradual shift to lower SFs with eccentricity (m_N_), SF bandwidth (b_N_) and amplitude (γ_N_). Consequently, attention operated identically on each texture stimulus, with the spatial spread of attention fixed across experiments (see Methods).

The model reproduced the central impairments, peripheral improvements and their variation with texture scale. For a fine-scale texture, the narrow SF profile yielded improvements within the parafovea (4-12°), impairments across a small range of central eccentricities (0-2°) and shifted peak performance toward the periphery (∼6°; Figure 5A). For the coarser texture, the same attention profile generated improvements in the periphery (8-22°), impairments within the parafovea (0-8°) and shifted peak performance farther toward the periphery (∼15°; Figure 5B).

**Figure 5.**
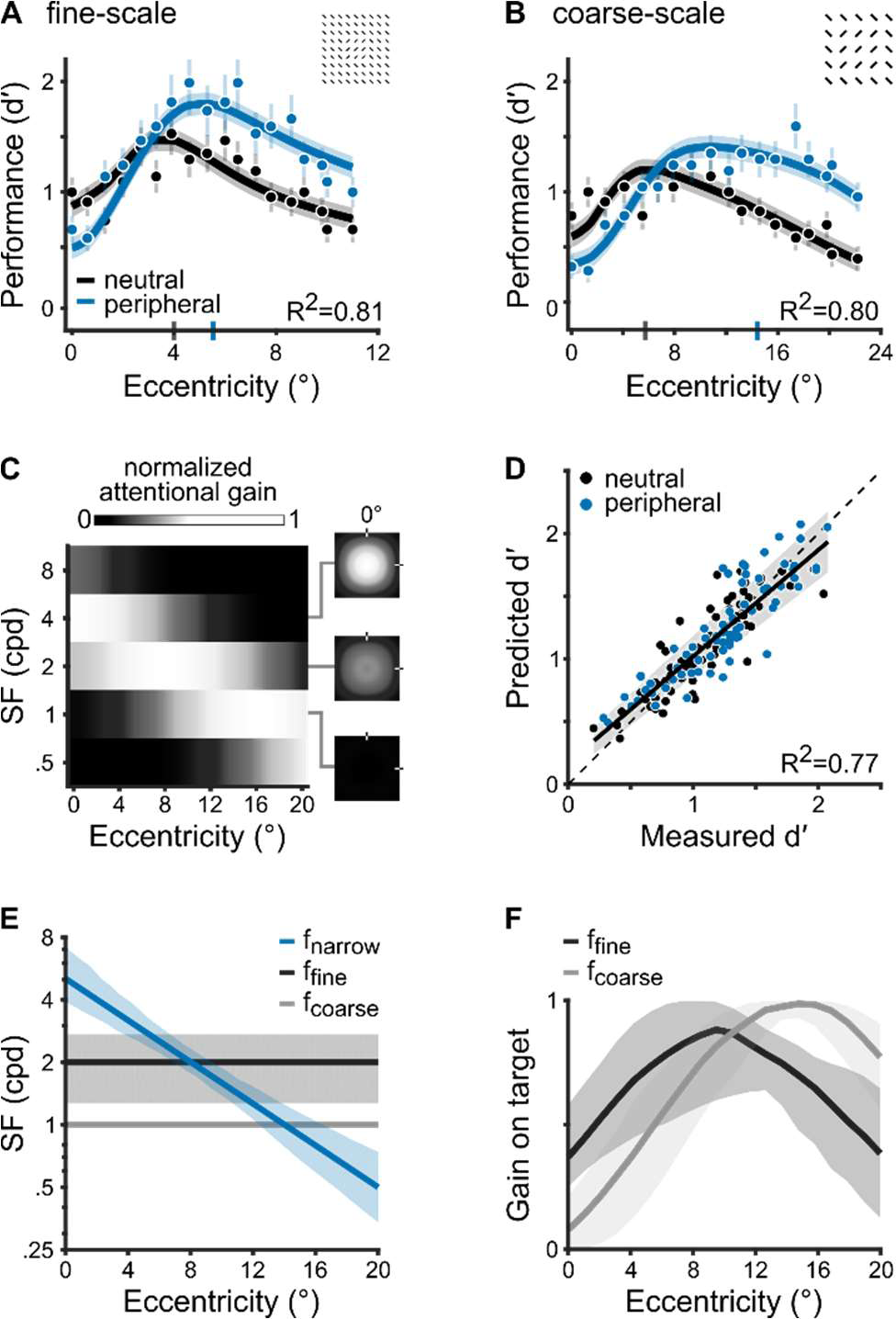
Narrow high-SF enhancement generates exogenous attention effects (A) Fit to Experiment 1 in Yeshurun & Carrasco, 1998 (27). The dots (n=36) depict group-average performance and error bars denote ±1 SEM. The solid lines and shaded regions indicate the median and 68% confidence intervals of the bootstrapped distribution of model fits. The vertical blue bar on the x-axis indicates the eccentricity of peak performance with peripheral cues. (B) Fit to Experiment 2 (n=36) in Yeshurun & Carrasco, 1998 (27). The model jointly fits performance on neutral and peripheral cue conditions with parameters shared among all six experiments, including the data shown in A. Visualization follows the conventions in A. (C) Best-fitting narrow gain profile. The matrix depicts attentional gain across eccentricity, normalized by the maximum and averaged across space and orientation. Matrix visualization and the panels on the right follow the conventions of Figure 4A. (D) Goodness-of-fit for neutral and peripheral-cued performance (n=146). Plotted as in Figure 4C. (E) SF preference of the narrow attentional gain profile (f_narrow_) and the SF that best distinguished fine-(f_fine_) and coarse-scale targets (f_coarse_). The solid lines and shaded areas indicate the median and 68% bootstrapped confidence interval. The shaded area for f_coase_ overlaps the solid line. (F) Normalized magnitude of attentional gain on the fine- and coarse-scale target SF across eccentricity (median and 68% confidence interval of bootstrapped distribution).

A gradual shift of attentional gain toward lower SFs (26) reproduced the transition from impairments to improvements across eccentricity (Figure 5C). At the fovea, attentional gain was centered on a SF (4 cpd) higher than those distinguishing the fine-(2 cpd, f_fine_) or coarse-scale (1 cpd, f_coarse_) textures. As a result, the population response shifted away from the target and impaired performance. With increasing eccentricity, attentional gain progressively overlapped the SF of each target, improving performance. Attention enhanced the fine-scale target SF within the parafovea (4-12°) then enhanced the coarse-scale target at farther eccentricities (8-22°). Overall, across the six experiments, the model explained 77% of the variance (95% bootstrapped CI = [49 82]; Figure 5D).

Attentional gain on SFs higher than the target yielded impairments at foveal locations. This pattern was consistent across all six experiments (Figure 5E). Consequently, the overlap between fine-(f_fine_) or coarse-scale (f_coarse_) targets and the SF tuning of attentional gain was minimal at the fovea and peaked in the periphery (Figure 5F). This mismatch between the SF tuning of attention (f_narrow_) and the target is suggested to be driven by exogenous attention operating above intrinsic SF preferences at each eccentricity (26). We corroborated this relation. We compared f_narrow_ to the model’s baseline SF tuning, indexed by the peak SF of the stimulus drive (f_stim_, Figure 2A). Consistent with empirical measurements, we found that the narrow SF profile preferred SFs higher than baseline tuning (Figure S2).

### Broad SF enhancements yield endogenous attention effects

The model predicted group-average data from neutral and central cueing conditions across four experiments (60 data points). Endogenous attention was modeled as a broad gain profile (Figure 2A, red) (26). 12 free parameters constrained model behavior (Table S4). Four parameters, shared among experiments, determined attentional gain–foveal SF preference (a_B_), the decline in SF preference with eccentricity (m_B_), SF bandwidth (b_B_) and amplitude (γ_B_).

The model reproduced improvements across eccentricity for both fine-(Figure 6A) and coarse-scale textures (Figure 6B). To generate these improvements, attentional gain encompassed the target SF for each texture scale (Figure 6C). Across all four experiments, the model explained 89% of the variance (95% bootstrapped CI [67 92]; Figure 6D).

**Figure 6.**
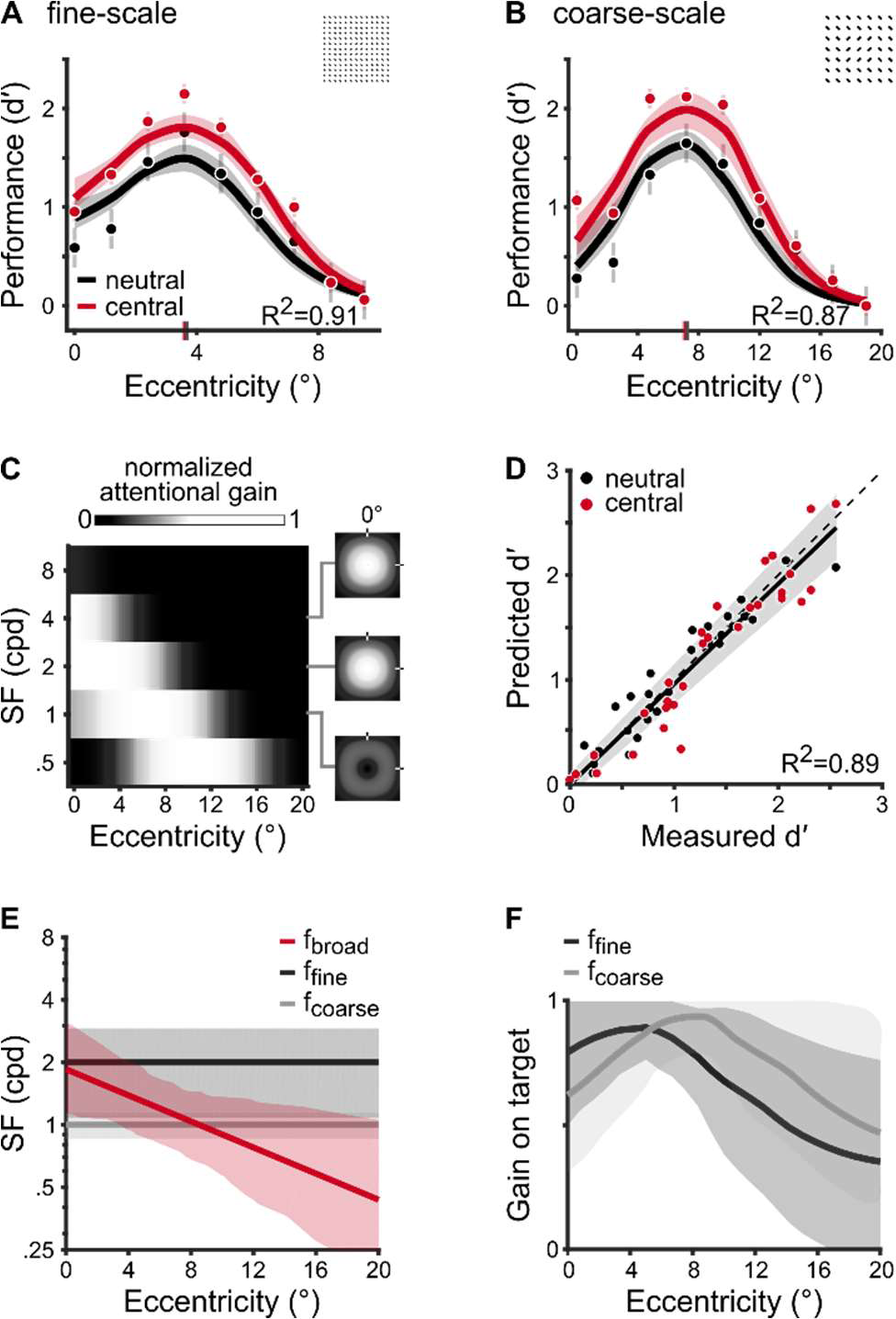
Broad SF enhancements yield endogenous attention effects (A) Fit to Experiment 3 in Yeshurun, Montagna & Carrasco, 2008 (32). The dots (n=18) depict group-average performance and error bars denote ±1 SEM. The solid lines and shaded regions indicate the median and 68% confidence intervals of the bootstrapped distribution of model fits. The vertical red bar on the x-axis indicates the eccentricity of peak performance with peripheral cues. (B) Fit to Experiment 4 (n=18) in Yeshurun, Montagna & Carrasco, 2008 (32). The model jointly fits performance on neutral and central cue conditions with parameters shared among all four experiments, including the data shown in A. Visualization follows the conventions in A. (C) Best-fitting broad gain profile. Plotted as in Figure 5C. (D) Goodness-of-fit for neutral and central-cued performance (n=60). Plotted as in Figure 4C. (E) SF preference of the broad attentional gain profile (f_broad_) and the SF that best distinguished fine-(f_fine_) and coarse-scale targets (f_coarse_). The solid lines and shaded areas indicate the median and 68% bootstrapped confidence interval. (F) Normalized magnitude of attentional gain on the fine- and coarse-scale target SF across eccentricity (median and 68% confidence interval of bootstrapped distribution).

Endogenous attention effects were reproduced by a broad SF attentional gain that was centered near the target SF across eccentricity (f_broad_ in Figure 6E). This contrasts with the narrow SF gain profile that modulated higher SFs at central locations to reproduce exogenous attention effects (Figure 5E). Although the center SF of attention declined with eccentricity, the modulation profile’s plateau ensured that it overlapped both fine- and coarse-scale target SFs across eccentricity (Figure 6F). Psychophysical measurements of attentional effects on contrast sensitivity (26) suggest that the SF range enhanced by endogenous attention is centered near those intrinsically preferred by an observer at each eccentricity. However, our model fits to texture segmentation experiments revealed that attentional gain enhanced lower SFs than baseline tuning (f_stim_) at central locations (Figure S3).

### Different SF gain profiles govern exogenous and endogenous attention effects

We directly assessed whether different SF gain profiles—narrow or broad—generate the effects of exogenous and endogenous attention. In addition, we compared the efficacy of SF-tuned gain against a model wherein the spatial extent of attention varied across experiments while the gain across SF was uniform. The spatial spread of attention is a key factor of the NMA (15), which posits that its extent relative to the stimulus size helps reconcile apparent discrepancies between each attention type’s effects on contrast sensitivity. These predictions have been empirically tested and confirmed (66). By comparing the narrow and broad SF models to the spatial extent model, we directly assessed the separate contributions of SF gain and the spatial spread of attention to segmentation performance (Figure 7).

**Figure 7.**
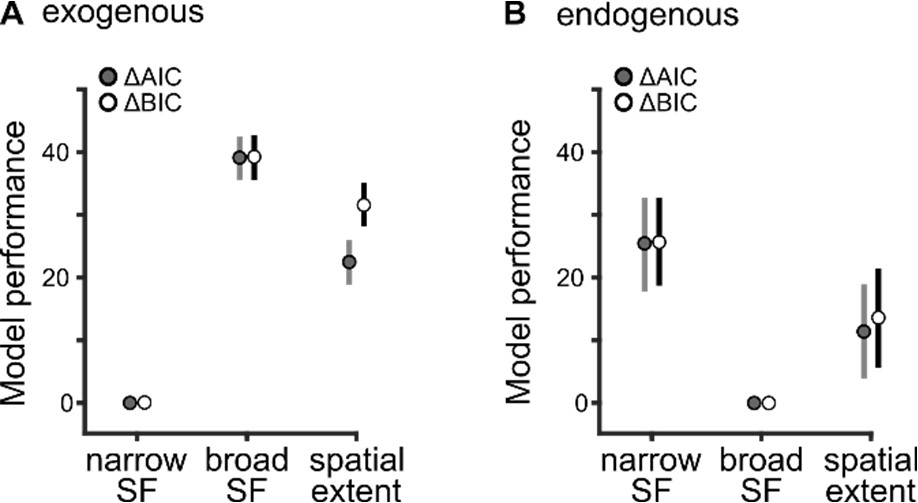
Different SF gain profiles govern exogenous and endogenous attention effects. AIC and BIC model comparisons for different regimes of attentional modulation for (A) exogenous attention and (B) endogenous attention. The dots and error bars represent the median and 95% confidence intervals of the bootstrap distributions. A parsimonious explanation for several experimental manipulations in texture segmentation Figure 8 depicts behavioral data for a variety of texture segmentation experiments. Whereas we focus on the impact of texture scale in Figure 5 and 6, the model is general. It jointly accounted for multiple target locations (vertical, Figure 8A; horizontal, Figure 8C-E; and intercardinal meridians, Figure 8F), behavioral tasks (orientation discrimination, Figure 8B) and attentional manipulations (cue size, Figure 8C). Although the model was fit using texture images with fixed positions and orientations (Figure 3), it behaved similarly for textures with randomly jittered elements (Figure S5). Overall, the proposed model provides a parsimonious explanation for and a quantitative match to segmentation performance (Figure 8).

Tuned SF gain modulation reproduced the effects of attention. The spatial extent alone was insufficient to capture the effects of either exogenous (ΔAIC: M=21.2, CI=[18.8 26.0]; ΔBIC: M=31.7, CI=[27.9 34.9]; Figure 7A) or endogenous attention (ΔAIC: M=11.4, CI=[3.9 18.9]; ΔBIC: M=13.5, CI=[5.7 20.8]; Figure 7B). For exogenous attention, the narrow profile outperformed the broad profile (ΔAIC: M=39.1, CI=[35.5 42.5]; ΔBIC: M=39.1, CI=[35.9 42.5]; Figure 7A). For endogenous attention, the broad profile outperformed the narrow profile (ΔAIC: M=25.4, CI=[17.8 32.7]; ΔBIC: M=25.5, CI=[18.0 32.7]; Figure 7B). Decrements in model performance manifested as an inability to capture impairments or improvements at eccentricities demarcating the CPD (Figure S4). Thus, these model comparisons substantiate psychophysical measurements (25, 26): exogenous and endogenous attention effects are best explained by different attentional gain profiles across SF.

### Model predictions generalize to basic visual tasks

To test whether this model generalizes to other basic visual tasks, we applied it to tasks mediated by acuity (6) and contrast sensitivity (26), with no additional model parameters (Figure 9). These studies separately manipulated exogenous and endogenous attention and highlight how attention effects depend on the stimulus and task. In the acuity task, observers discriminated the location of a small gap (<1°) in a Landolt square (Figure S6A) whereas contrast sensitivity was measured with gratings in an orientation discrimination task (Figure S7A).

The model reproduced the improvements to acuity and contrast sensitivity for each attention type. On the one hand, both exogenous and endogenous attention improve acuity similarly (6). Model simulations yielded consistent visual acuity improvements for both exogenous (Figure 9A) and endogenous (Figure 9B) attention, despite different SF gain profiles underlying each attention type. On the other hand, each type of attention alters contrast sensitivity across SF differently (26). Model simulations captured the differences between exogenous (Figure 9C) and endogenous attention (Figure 9D). The model reproduced the narrow SF bandwidth of exogenous attention that is centered on SFs higher than baseline tuning preferences (Figure S7D). It also captured the broad SF modulation by endogenous attention that spanned SFs above and below baseline tuning (Figure S7E). Attention effects derived from our observer model closely matched descriptive fits to the data from (26) (Figure 9C-D).

**Figure 8.**
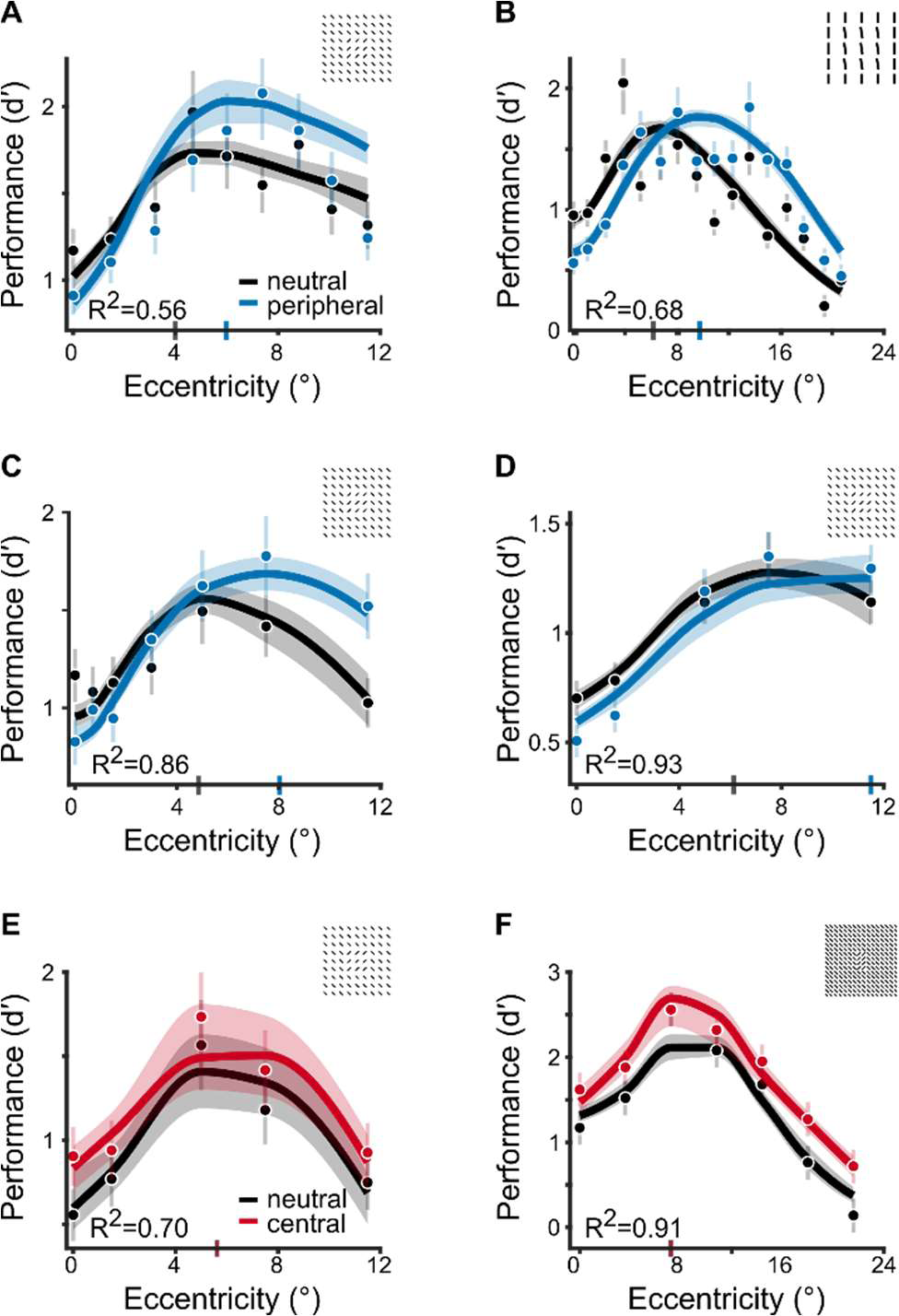
A parsimonious explanation for several experimental manipulations in texture segmentation (A-D) Narrow SF gain profile fit to exogenous attention experiments. The model jointly fits these data and those displayed in Figure 5, with parameters shared among all six experiments. Insets in each panel depict the same textures displayed in Figures 3C-F, respectively. (E-F) Broad SF gain profile fit to endogenous attention experiments. The model jointly fits these data and those displayed in Figure 6, with parameters shared among all four experiments. Insets in each panel depict the same textures displayed in Figure 3G and 3J, respectively. The dots and error bars depict group-average and ±1 SEM. The solid lines and shaded regions depict the median and 68% confidence intervals of the bootstrapped distribution of model fits.

**Figure 9.**
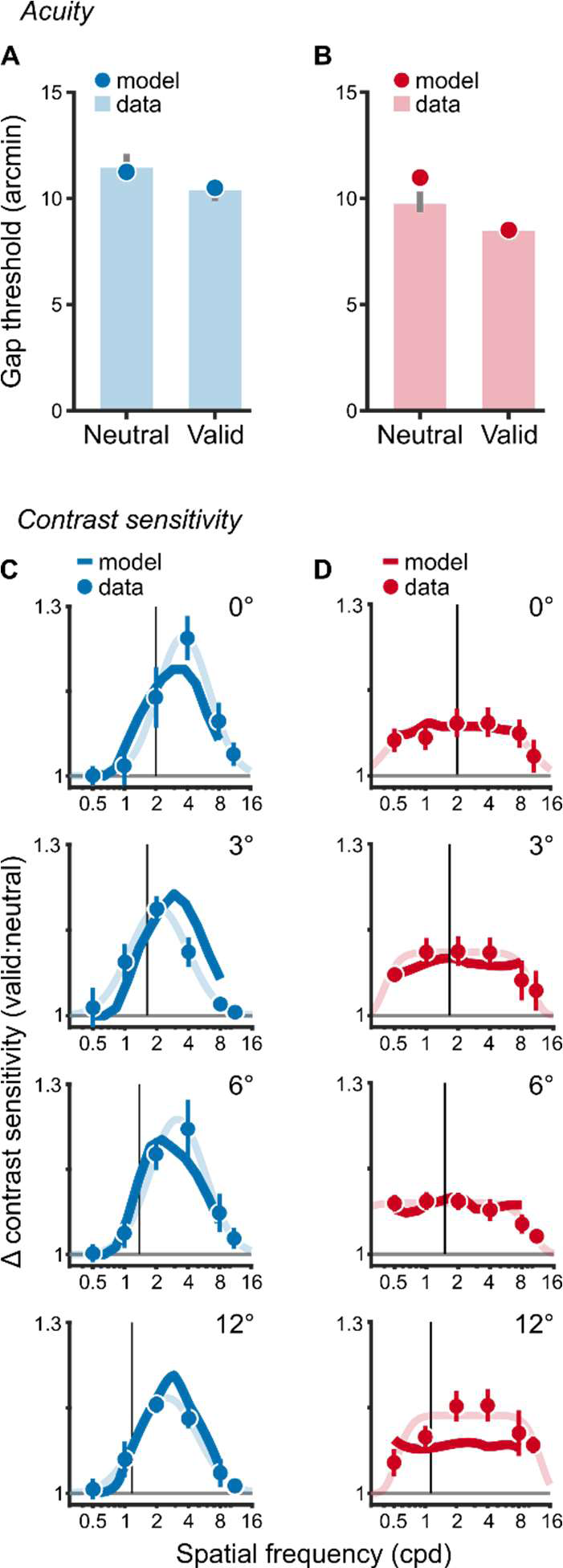
Model predictions generalize to other basic visual tasks. The effects of (A) exogenous and (B) endogenous attention on gap discrimination thresholds in an acuity task. Data from (6). Lower thresholds indicate higher acuity. Bars depict group-average thresholds in neutral and valid cueing conditions. Error bars are ±1 SEM. Dots depict model-derived gap thresholds for the acuity task. (C) Exogenous and (D) endogenous attention effects on contrast sensitivity across SF and eccentricity, quantified as the ratio between valid and neutral contrast sensitivity. Data from (26). Values above 1 indicate an attentional enhancement of contrast sensitivity. The dots and error bars depict the group-average and ±1 SEM. The vertical black lines show baseline SF preferences measured in the neural condition (Figure S7). The solid colored lines show model fits to the data whereas lightly shaded lines are descriptive fits to the data from (26). In all panels, the narrow SF profile was fit to exogenous attention effects whereas the broad SF profile was fit to endogenous attention effects.

The attention parameters were consistent across tasks (Table S6). The SF bandwidth of endogenous attentional gain consistently spanned a larger range than exogenous attention (Table S6, SF bw). Moreover, the rate at which SF selectivity declined with eccentricity also differed. The peak SF decreased with eccentricity (Table S6, SF slope), but less so for exogenous than endogenous attention, indicating that exogenous attention consistently enhanced SFs higher than the peak SF of the stimulus drive (Figure S2). Lastly, we observed tradeoffs between the amplitude and spatial spread of attention (Table S6). In the acuity task, the amplitude was large (>8) and the spatial spread was narrower (0.6°) than the stimulus (1°), whereas in contrast sensitivity, the amplitude was lower (<1.5) and the spatial spread was broader (>5°) than the stimulus (4°). Texture segmentation yielded intermediate values wherein the amplitude was ∼4 for a fixed spread of 4°. Independent of attentional effects, differences in the experimental protocol and stimuli used across experiments resulted in subtle differences in the best-fitting model parameters for contrast gain and the stimulus drive. Importantly, similar attention parameters reproduce endogenous and exogenous attention effects in a variety of visual tasks.

## DISCUSSION

We used texture segmentation as a model system to dissociate endogenous and exogenous attention. To this end, we developed an image-computable model that reproduces human segmentation performance and the modulations by each attention type. This model links neural computations to three visual phenomena. (i) Divisive normalization and spatial summation mediate the CPD (27-34, 43-46). (ii) Narrow high-SF enhancement drives exogenous attentional effects (27–32, 34). (iii) Broad SF gain drives endogenous attentional modulations (32–34).

Normalization models of attention have described how spatial attention affects neural responses and behavior (e.g., (14, 15, 17)). Our model adopts the same algorithm specified by the Reynolds-Heeger normalization model of attention (15) (NMA)—attentional gain modulates the stimulus drive before divisive normalization. Predictions by NMA have been empirically confirmed with psychophysical experiments (66). These experiments equated seemingly distinct effects of endogenous and exogenous attention on contrast sensitivity by manipulating and accounting for the spatial extent of attention.

Here, we demonstrate a critical limitation of extant models of attention. Their predictions do not extend to the differential effects on spatial resolution and do not explain the dissociation between endogenous and exogenous attention. Although the spatial extent of attention is critical for explaining effects on contrast sensitivity, our model comparisons demonstrate that it is not vital for reproducing attention effects on texture segmentation (‘spatial extent’ model in Figure 7 and Figure S4). These results corroborate empirical evidence that manipulating the spread of attention during texture segmentation does not yield shifts between the typical effects of endogenous and exogenous attention (31).

To capture the effects of attention on texture segmentation we implemented: (i) Eccentricity-dependent and SF-tuned multiplicative gains that emulate neural (54) and psychophysical (53) SF selectivity. (ii) Spatial summation, which emphasizes textural contours (39, 41, 47). (iii) Distinct SF gain profiles for endogenous and exogenous attention (25, 26) that scale responses prior to normalization (15), thereby adjusting the balance between fine and coarse-scale visual sensitivity. The model’s distinct SF profiles instantiate a computational dissociation between each attention type that substantiates their differential impact on sensory processing.

The necessity for different SF profiles is supported by empirical evidence (25, 26) and provides insights toward the distinct roles of endogenous and exogenous attention in guiding visual behavior. Previous models (e.g. (14, 15, 17)) demonstrate that both forms of attention improve low-level visual processes that encode elementary features (e.g., contrast, orientation, motion). Here, we show that attention differentially interacts with normalization to shape the competition inherent in mid-level processes such as texture segmentation. Exogenous attention preferentially enhances a narrow range of high SFs. Consequently, its effects prioritize fine-grained visual details at the expense of competing coarse-scale features within a stimulus. In contrast, endogenous attention consistently improves mid-level processing by broadly enhancing sensory encoding across fine and coarse spatial scales. The computations underlying mid-level processing bridge the gap between sensory encoding and object recognition (39–42). Therefore, the distinct impact by each type of attention and their computational differences at this processing stage have broad implications for natural visual behavior.

The model provides a computational framework for understanding the mechanisms underlying established effects of exogenous attention on spatial resolution (27–34) (reviews (1–3)). Previous studies offered qualitative descriptions that exogenous attention automatically increases spatial resolution (27–32, 34) (reviews (1–3)) with concomitant costs in temporal resolution (22) attributed to an engagement of parvocellular neurons (22, 67). Here, we develop an observer model that anchors these qualitative descriptions onto established neural computations. In doing so, we corroborate previous psychophysical experiments that found a similar high-SF preference of exogenous attention (25, 26, 30, 68), specify how attentional gain changes across the visual field and demonstrate its computational validity for explaining effects on perception.

We also provide converging evidence that exogenous attention alters perception inflexibly. By comparing the model’s exogenous attentional gain on textures to empirical measurements made with gratings (26), we found that it consistently operates above intrinsic (i.e., baseline) SF preferences despite large differences in stimuli (Figure S2). These findings suggest that in addition to exogenous attentional effects being invariant to cue validity (8) and sometimes detrimental to perception (27–32, 34), its operating range across SF is also invariant to the type of stimulus being attended.

The model provides insights on the mechanisms underlying endogenous attention effects on spatial resolution. Previous research has established that endogenous attention modulates texture segmentation (18, 32–34, 69) and its impact has been described as an optimization of spatial resolution (reviews (1–3)). We propose that a broad SF gain control mechanism yields these perceptual improvements. Our proposal complements previous reports that endogenous attention uniformly excludes noise across SF (70), but seemingly conflicts with an earlier explanation that endogenous attention suppresses sensitivity to high SFs to improve texture segmentation (33). However, suppressed high-SF sensitivity at foveal locations would decrease cross-frequency suppression (59, 61) and result in an effective dominance of lower SFs, which is compatible with our findings (Figure S3).

Moreover, we provide converging evidence of the flexibility of endogenous attention. We found that the model’s SF preference during texture segmentation differed from those measured with gratings (26). This discrepancy suggests that the impact of endogenous attention depends on the properties of the attended stimulus and the nature of the task, consistent with the notion of a flexible endogenous attentional mechanism (8, 32–34).

The effects of attention depend on divisive normalization. Without normalization, we could not reliably capture the CPD, which served as the foundation of our analyses. Previous studies demonstrate that when the pool of SFs contributing to normalization is restricted, the CPD is attenuated (30, 33, 44). However, existing models of the CPD (46, 48, 49) relate the phenomenon solely to an increase in receptive field size with eccentricity. Our model directly links the summation area of receptive fields to their SF tuning. Consequently, the dominant summation area increases with eccentricity as SF preferences decrease. Despite implementing an increase in receptive field size, we could not capture the CPD without accounting for the surrounding context via normalization.

Additionally, we demonstrate that spatial constraints mediate the CPD independently from limitations in temporal processing across eccentricity. The proposal that the CPD may result from slow information accrual at the fovea, which yields poor performance particularly when a backward mask limits processing time (43), has been criticized (45, 46, 71). We note that our model accounts equally well for the findings of texture segmentation studies regardless of whether they contained or omitted a mask, which minimized temporal contributions to task performance (Table S5).

Importantly, both endogenous and exogenous attention speed information accrual (72) across the visual field (73, 74) and across different levels of cue validity (8). Thus, effects of attention on temporal processing would predict similar improvements by each attention type on the CPD, a prediction clearly contradicted by the modeled studies here (27, 29–33).

The computations implemented in the model are based on the known properties of the human and non-human primate visual system. The stimulus drive simulates bottom-up responses of phase-invariant complex cells in V1 (56) that vary with SF and eccentricity (53, 54). The model’s response to texture is generated through pooling bottom-up inputs, consistent with the gradual emergence of texture selectivity along the visual hierarchy (75–77).

Exogenous attentional gain in the model result in changes to texture sensitivity; however, little is known about the neural underpinnings of these effects. There are sparse demonstrations of exogenous attentional modulations in visuo-occipital areas and beyond (20, 21, 78–80). Transcranial magnetic stimulation of early visual cortex reveals that its activity plays a key role in the generation of exogenous attention effects (81). However, future studies are required to determine how the SF gain modulation we report manifests in neural populations.

In contrast, it is established that endogenous attention modulates cortical responses (1, 2, 13, 18, 20, 21, 36–38, 82, 83). During texture segmentation tasks, endogenous attention selectively enhances V1 and V4 responses to the embedded figure, suggesting that attention spreads across the target object to facilitate its segmentation (18). Our model provides complementary evidence that endogenous attention optimizes SF sensitivity to improve segmentation across texture scale. Yet, it is unclear how neural activity generates these SF modulations. Neuroimaging (37, 38) and electrophysiological (13, 36) recordings demonstrate that spatial tuning profiles are altered by endogenous attention. Such changes are consistent with, but not necessary for, the modulations of spatial resolution we report.

Few computational models have implemented possible ways in which attention alters spatial resolution. Some have proposed that attention modifies how finely a spatial region is analyzed. Such changes are either driven by an attention field that adjusts the spatial profile of receptive fields (13) or by attracting receptive fields toward and contracting them around the attended location (19). Other models suggest an attentional prioritization that selectively tunes responses for a given spatial location and attenuates responses to surrounding regions (12, 16). However, these models neither account for differences across eccentricity nor explain attentional shifts toward fine or coarse spatial scales. Critically, these models do not distinguish between endogenous and exogenous attention. In contrast to these previous models, we do not propose any modifications to the structure of receptive fields. Instead, we attribute changes in spatial resolution to modulations of SF, a fundamental dimension of early visual processing.

The fact that our model operates on arbitrary images facilitates its generalization to other visual stimuli and tasks. We show that the model reproduces the differential endogenous and exogenous attention effects on contrast sensitivity (Figure 9C-D). Notably, the model recreates behavior in visual acuity tasks where the improvements by each attention type are similar (Figure 9A-B).

Unlike texture segmentation, acuity tasks always benefit from heightened spatial resolution, which obscures differences between these two attention types. Recent studies that compared both attention types head-to-head with the same observers, stimuli and task found that they produced similar behavioral effects but modulated neural activity differently in the temporo-parietal junction (20) and occipital cortex (21). Our model is consistent with these findings and highlights that differences in the underlying computations can yield similar perceptual effects between endogenous and exogenous attention depending on the stimulus and task.

Future work may extend the model to other visual phenomena. For instance, it could capture the differential effects by each attention type on second-order texture perception (28, 34), second-order texture contrast sensitivity (24) and temporal resolution (22, 23, 67). Lastly, it is unknown how interactions between both forms of attention may affect mid-level processes like texture segmentation. Endogenous attention attenuates the transient effects of exogenous attention on stimulus discriminability when both are deployed concurrently (84). Therefore, it is possible that endogenous attentional benefits will outweigh the costs induced by exogenous attention when both are deployed simultaneously during texture segmentation. Although the experimental designs of the studies we have modeled cannot address this open question, our model framework may facilitate predictions of the perceptual consequences when both forms of attention are deployed.

In conclusion, we reproduce signatures of texture segmentation (27-34, 43-46) and characterize the contributions of attention to a process commonly considered ‘preattentive’ (39, 41, 42, 44–49). Moreover, we reveal the neural computations that underlie how attention modifies spatial resolution (1–3). Attention scales sensitivity to high and/or low SFs, adjusting the balance between fine and coarse-scale spatial resolution. Exogenous attention preferentially enhances fine details whereas endogenous attention uniformly enhances fine and coarse features to optimize task performance. Because the model distinguishes between endogenous and exogenous attention, varies with stimulus eccentricity, flexibly implements psychophysical tasks and operates on arbitrary grayscale images, it provides a general-purpose tool for assessing theories of vision and attention across the visual field.

## METHODS

### Model

We developed an observer model that simulates the response of a collection of receptive fields (RFs) each narrowly tuned to spatial position (x,y), orientation (*θ*) and SF (f). Responses varied with eccentricity (*α*). The population response (R) is generated by four components: the stimulus drive (E), attentional gain (A), suppressive drive (S and σ), and spatial summation (F), where * represents convolution:

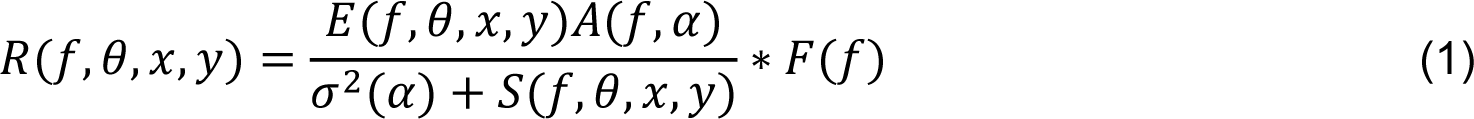

All model parameters are given in Table S1.

### Stimulus drive

The stimulus drive characterizes responses of linear RFs in the absence of suppression, attention and spatial summation. A steerable pyramid (55) decomposed stimulus images into several SF and orientation subbands, defined by weighted sums of the image (i.e., linear filters). Weights were parameterized by raised-cosine functions that evenly tiled SFs, orientations and positions.

The number of SF and orientation subbands are parameters that can be flexibly chosen. We used a set of 30 subbands comprising five SF bands and six orientation bands. The size of the stimulus image and the subband bandwidth determine the total number of SF subbands. In our simulations, images were 160 x 160 pixels (see Stimulus generation) and SF bandwidth (i.e., full-width at half-maximum, FWHM) was 1 octave, which allowed for five different SF subbands. The chosen bandwidth is comparable to empirical tuning curves measured in primate electrophysiological recordings (85) and human psychophysical (53) measurements. The FWHM orientation bandwidth (60°) is comparable to physiological tuning curves measured in primates (86). Using narrower (30°) or wider (90°) bandwidths yielded similar results supporting the same conclusions.

The pyramid includes RFs in quadrature phase. We computed a ‘contrast energy’ response (56), (i.e., the sum of squared responses across phase) which depends on the local spectral energy at each SF, orientation and position in the image. Contrast energy is fundamental to texture perception models (39, 41, 47) and we denote it as C(f,*θ*,x,y). SF gain. Human (26, 53, 87) and non-human primate (54) contrast sensitivity is narrowly tuned to SF. SF tuning shifts from high to low SFs with eccentricity. To model this behavior, contrast energy was multiplied point-by-point by a SF gain function, T, defined by a log-parabola (88, 89):

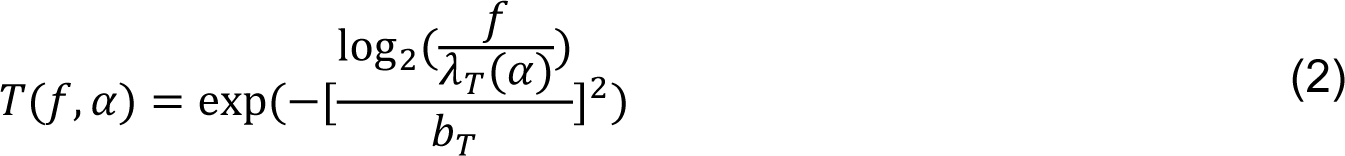

where *α*denotes the eccentricity of a RF and b_T_ determines the function’s SF bandwidth. The preferred SF (*λ*T) at a given eccentricity is given by:

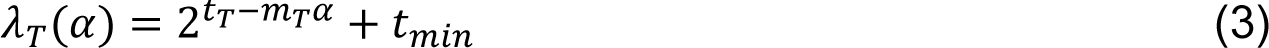

SF preferences converge onto a single value in the far periphery, t_min_(87). The preferred SF at the fovea is given by 2 + t_min_and progressively shifts towards t_min_at the rate m_1_. Whereas t_1_ varied during simulations (see Table S1-S4), t_min_was fixed at 0.5 cpd because texture stimuli produced minimal contrast energy below that SF subband. Allowing t_min_to vary yielded similar results supporting the same conclusions.

In sum, the stimulus drive (E) characterizes the contrast energy responses that vary with SF and eccentricity, computed as:

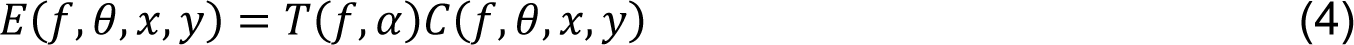

### Attentional gain

Attention is implemented as an attentional gain field, A, that multiplies the stimulus drive point-by-point as in the Reynolds-Heeger normalization model of attention (15). Attentional gain was uniform across orientation. Across SF and position, gain was distributed according to cosine window functions, w:

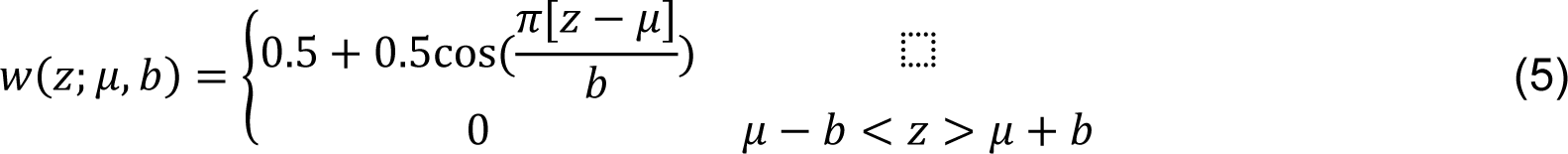

where *μ*defined its center and b defined its FWHM. The units of *μ* and z depended on the dimension: for SF each variable was in units of log_2_-transformed cycles per degree and for position they were in units of degrees of visual angle. The window was defined on a logarithmic axis for SF but on a linear axis for position. SF and spatial position functions were multiplied, point-by-point, to characterize the full distribution of attentional gain.

Spatial spread. Attentional gain was centered on the target location. In our simulations, the target fell along the horizontal meridian at eccentricity *α*targ (see Stimulus generation). The product of two cosine functions (w, equation 5) defined the spread of attention: one varied as a function of x and another as a function of y, each with an identical width b_pos_. Widths did not vary across eccentricity. A_pos_ defined the spatial spread of attention:

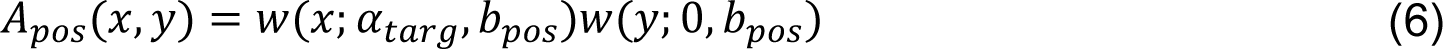

The precise spatial spread of attention is controversial (90) and can change based on task demands (66, 91). Critically, it has not been explicitly manipulated during texture segmentation tasks by varying the target’s spatial uncertainty. Such a protocol has been used to test predictions of the NMA and has been demonstrated to adjust the size of the attention field (66). Instead, a previous study (31) measured exogenous attention effects while manipulating the size of a peripheral pre-cue. The authors found that exogenous attention altered performance as long as the cue was the same or smaller than the target size. In our simulations, the spread of attention was fixed at a FWHM of 4° (Table S1) because it encompassed the largest target size used to constrain model parameters (Table S5). As a result, the spatial extent of attention was identical across eccentricity and experiments. Similar results were observed when the spread was fixed at 2° and 3°. However, in the model variant wherein the spatial extent could change (see Model alternatives), the FWHM of attentional spread (b_pos_) was free to vary between experiments.

SF gain profile. We implemented two gain profiles: narrow and broad (26).

Narrow profile. In the narrow model (AN), attentional gain was bandpass across SF. Attentional gain peaked at a given SF, *λ*N, and fell gradually toward neighboring frequencies within its bandwidth, b_N_, characterized by a cosine function:

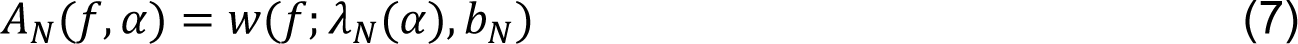

The center SF of attentional gain profiles (*λ*N for narrow, *λ*B for broad) varied with eccentricity:

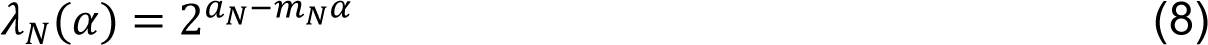

where a_N_ (or a_B_ for the broad profile) defined the center frequency at the fovea, which gradually changed with eccentricity at the rate m_N_ (m_B_ for broad).

Broad profile. The broad profile (A_B_) implemented broadband attentional gain, characterized by the sum of three overlapping cosine functions:

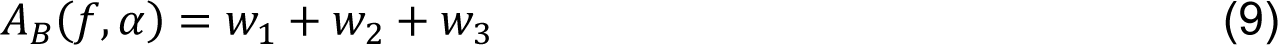

where 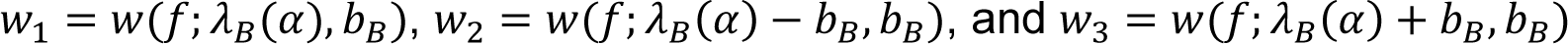. The bandwidth of each function was given by b_B_. Relative to the center SF, *λ*B, the adjacent functions were centered ±b_B_ apart, ensuring that their sum yielded a plateau spanning b_B_ octaves and a FWHM of 1.5b_B_.

In sum, attentional gain multiplicatively scaled the stimulus drive uniformly across orientation, but differently across SF and eccentricity given by:

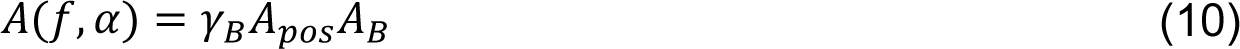

where A_pos_ and A_B_ (or A_N_) were four-dimensional matrices characterizing attentional gain across position, SF and orientation. *γΒ* (or *γN*) defined attentional amplitude. To simulate the neutral cueing condition, amplitude was set to 1. In addition, to assess the explanatory power of the spatial spread of attention (see Model alternatives), A_B_ (or A_N_) were set to 1 and only *γ* and A_pos_ varied.

### Suppressive drive

The suppressive drive comprised contextual modulation, computed through pooling the attention-scaled stimulus drive (15) across nearby positions, all orientations and neighboring SFs. This pooling procedure implemented lateral interactions between RFs and was computed via convolution (15). Convolution kernels were cosine window functions (w, equation 5).

The bandwidth of the SF kernel, δ_f_, equaled 1 octave:

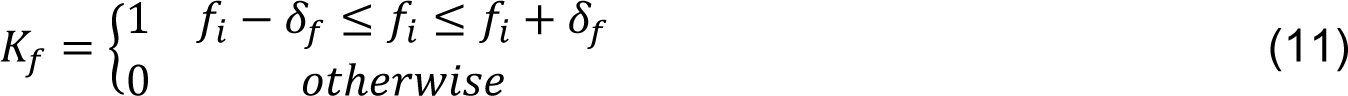

where fi denotes the center SF of a subband. This kernel summed contrast energy within and ±1 octave around each SF subband.

The bandwidth of the orientation kernel, δθ, equaled 180°, which encompassed all orientation subbands:

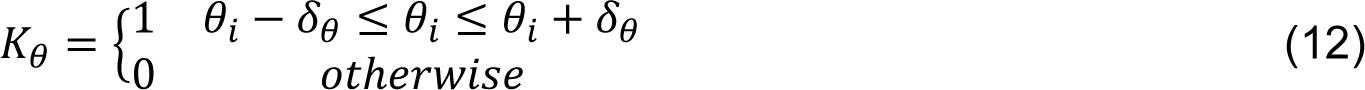

where θi denotes the center orientation of a steerable pyramid subband. This kernel summed contrast energy across all orientations.

Spatial position kernels were determined by multiplying two cosine windows:

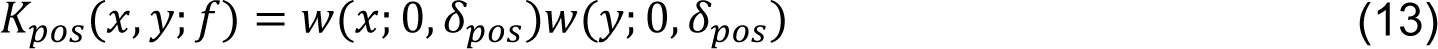

One window varied across x, another across y and their centers traversed across the image during convolution. The two-dimensional kernel summed to unity, which computed the average energy within the pooled area. Kernel width, δ_pos_, equaled and was inversely proportional to subband SF *f* and yielded two-dimensional spatial kernels, K_pos_. Kernel widths were identical across eccentricity. Contextual modulation was characterized via separable convolution:

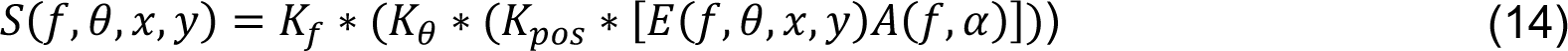

where * denotes convolution of the suppression kernels, K. Suppression magnitude was adjusted across eccentricity by σ^2^, which controlled the level of contrast at which neural responses reached half-maximum and is referred to as contrast gain. Contrast gain was implemented as an exponential function across eccentricity (26, 87):

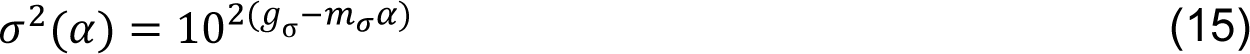

where g_σ_ and m_σ_ are free parameters that determine contrast gain at the fovea and the rate at which it varies with eccentricity, respectively.

### Spatial summation

Following divisive normalization, responses were weighted and summed across space, within each SF and orientation subband. Summation was accomplished via convolution by cosine windows, F, computed using equation (13). The width of each filter scaled with SF: narrow (wide) regions of space were pooled for high (low) SFs (39) and did not vary with eccentricity.

### Decision mechanism

We used signal detection theory to relate population responses to behavioral performance (d′). The available signal s was computed as the Euclidean norm of the difference between target-present (r_t_) and target-absent (r_n_) neural population responses: S = ||*r_t_ − r_n_*||. Performance on a discrimination task is proportional to the neural responses given the assumption of additive, independent and identically distributed (IID) noise. An alternative model with Poisson noise and a maximum-likelihood decision rule yields the same linkage between neural response and behavioral performance (92, 93). The signal and noise magnitude (σ_n_) defined behavioral performance d^’^ = 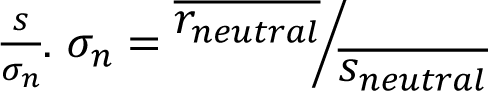 where 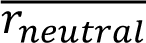 denotes the observed neutral performance averaged across eccentricity and S_1_ denotes the eccentricity-average of the signal. This ratio scaled the model’s predicted behavioral performance to match the observed data.

### Model fitting

Models were optimized by minimizing the residual sum of squared error between model and behavioral d′ using Bayesian adaptive direct search (BADS (94)). When applicable, performance data for a psychophysical experiment were converted from proportion correct, p, to d′ with the assumption of no interval bias (95): d^’^ = √2Z(p) where z denotes the inverse normal distribution. Although performance on 2IFC tasks can exhibit biases between intervals (96), our conversion algorithm operated uniformly across eccentricity, which preserved the performance variation (i.e., the CPD) critical for the goals of this study.

### Optimization strategy

#### Central performance drop

We fit the model jointly to performance on the neutral condition of all 10 texture segmentation experiments (103 data points). Peripheral and central cueing conditions (for exogenous and endogenous attentional conditions, respectively) were excluded to isolate the CPD. 15 free parameters fit all 103 data points (Table S2). Ten separate free parameters independently controlled the minimum contrast gain at the fovea (g*σ*; equation 15) for each of the 10 experiments. Sensitivity to contrast and SF varies for stimuli placed at isoeccentric locations around the visual field; it is higher at the horizontal meridian and decreases gradually towards the vertical meridian (97–100). Whereas 5 out of 6 exogenous attention experiments used targets placed on the horizontal meridian, 3 out of 4 endogenous attention experiments used targets presented along the intercardinal meridians (Table S5). Because SF selectivity depends on stimulus polar angle, two parameters separately determined the highest preferred SF (t_T_; equation 3)—one shared among exogenous attention experiments and the other shared among endogenous attention experiments. Alternatively, we could have fit separate parameters for horizontal, vertical and intercardinal meridians. However, this approach would have added a third free parameter, reducing the parsimony of the model. The configuration we used yielded reasonably good fits.

The remaining three parameters were shared among all experiments. Each controlled the bandwidth (b_T_; equation 2) of the tuning function T, the gradual shift toward lower SFs with eccentricity (m_T_; equation 3) and the increase in contrast gain across eccentricity (mσ; equation 15).

### Attentional modulation

To generate the effects of attention, the model was fit separately to exogenous and endogenous attention experiments. We jointly fit the model to neutral and valid conditions of each experiment.

Exogenous attention. All six exogenous attention experiments were fit jointly (146 data points) with 14 free parameters (Table S3). Minimum contrast gain at the fovea (g*σ*; equation 15) was determined independently for each of six experiments, yielding six free parameters. The remaining eight parameters were shared among all experiments. Four determined the stimulus drive: its bandwidth (b_T_; equation 2), the highest preferred SF at the fovea (t_T_; equation 3), the shift to lower SFs with eccentricity (m_T_; equation 3) and the slope of contrast gain across eccentricity (mσ; equation 15). The remaining four controlled the narrow SF attentional gain profile, specifically its bandwidth (b_N_; equation 7), center SF (λ_N_; equation 8), the shift to lower SFs with eccentricity (m_N_; equation 8), and its amplitude (γ_N_; equation 10).

Endogenous attention. All four endogenous attention experiments were fit jointly (60 data points) with 12 free parameters (Table S4). Minimum contrast gain at the fovea (gσ; equation 15) was determined independently for each experiment, which yielded four free parameters. The remaining eight parameters were shared among experiments, as described above for exogenous attention.

### Model alternatives

To assess whether contextual modulation and spatial summation are critical for the CPD, we implemented five model variants. Individual components of the suppressive drive were iteratively removed: cross-orientation suppression (‘-θ‘), cross-frequency suppression (‘-f’), surround suppression (‘-x,y’) and all components simultaneously (‘-all’). In a separate variant, spatial summation was removed (‘-sum’). We fit each variant separately to neutral performance data from all ten psychophysical experiments using the configuration described in Optimization strategy, Central performance drop.

In the ‘-all’ model, each RF was suppressed by its own response, simulating an extremely narrow suppressive pool. Specifically, the extent of suppressive pools (δ_f_, δ_θ_, δ_pos_; equations 11-13) were set to 0. As a result, the contributions of surround, cross-orientation and cross-frequency suppression were absent. The other contextual modulation variants only had a single parameter set to 0 (e.g., δf for cross-frequency suppression). The ‘-sum’ variant removed spatial summation (i.e., F in equation 1) from the model.

We additionally compared the efficacy of each attentional gain profile across SF—narrow or broad—in generating the effects of exogenous and endogenous attention by fitting each profile to exogenous and endogenous attention experiments. To assess the explanatory power of the spatial extent of attention, a third model was compared in which the spatial spread of attention (b_pos_, equation 6) varied between experiments and the gain across SF was uniform. Each model fit followed the configurations described in Parameter configuration, Attentional modulation.

### Model comparisons

We compared models using AIC (64) and BIC (65). The difference in AIC/BIC values between model variants indexed model performance. ‘-θ‘, ‘-f’, ‘-x,y’, ‘-all’ and ‘-sum’ models were compared to the full model. Additionally, narrow and broad SF gain profiles as well as the spatial extent model were compared.

### Stimulus generation

Target-present and target-absent textures were re-created to match the stimulus parameters used in each psychophysical study (Table S5). For all stimuli, each pixel subtended 0.03125° (i.e., 32 pixels/°), roughly matching the spatial resolution of a 1280 × 960 monitor display placed 57 cm away from the observer.

The full texture stimulus used in each experiment typically spanned the entire display. We generated 5°-wide square cutouts of the texture stimulus, centered on the target location. Because the model implemented visual sensitivity that varied with eccentricity, but was uniform at isoeccentric locations, all targets were assumed to be presented along the horizontal eccentricity for simplicity (as in equation 6).

Each texture array was composed of lines oriented 135°. The target comprised a patch of lines that were oriented 45°. One study was an exception (30) because the texture array comprised vertical lines (0°) and the target patch contained lines tilted ±8° (Figure 3D). In this study, observers’ performed an orientation discrimination task by reporting the orientation of the target presented on each trial. To simulate orientation discrimination performance, the target-present and target-absent stimuli always contained a patch but their orientation differed.

To avoid edge artifacts, texture stimuli were windowed by the sum of three cosine window functions (as in equation 8) centered on the target that produced a uniform plateau covering the central 3.75 deg and fell off with cosine edges. Pixel intensities in each stimulus were constrained between 0 and 1.

Textures used to fit the model were generated without spatial or orientation jittering. In additional simulations, the stimuli of two representative experiments were jittered. The stimuli for Experiment 1 in (27) were spatially jittered (0.3 deg jitter), and the stimuli in Experiment 4 in (32), were jittered spatially (0.34 deg jitter) and in orientation (55° bandwidth). Jitter parameters were compatible with those specified in each study.

### Resampling procedures

We obtained confidence intervals on the parameter estimates, model predictions and AIC/BIC values by bootstrapping the data and refitting the model 100 times per configuration (Optimization strategy) and for each model variant (Model alternatives). Bootstrap samples were generated by drawing and fitting random samples from Gaussian distributions centered on group-average performance at a given eccentricity, with the SEM for each study defining the distribution’s width.

To generate bootstrap samples for simulations with jittered texture stimuli, the model was first fit to the data for each experiment using a non-jittered texture. Then, the model parameters were fixed and jittered stimuli were input to the model. This procedure allowed us to assess how a fixed model behaved with variable texture inputs. One hundred unique jittered stimuli were presented to the model.

### Cross-validation procedure

To characterize how the operating range of exogenous and endogenous attention varied with eccentricity, relative to baseline tuning preferences (Figure S2-S3), we fit polynomials to empirical measurements made by (26). Leave-one-subject-out cross-validation determined the best-fitting polynomial order. Specifically, the ratio, in octaves, between the peak SF of the neutral contrast sensitivity function and the preferred SF of attentional modulation were computed for individual observers. Eccentricities were aggregated between each of the two experiments conducted. The ratio for one observer was set aside, and the remaining were averaged. Zero to second-order polynomials were fit to the group-average ratio across eccentricities. The sum of squared error to the left-out data point indexed cross-validation error. This process was iterated until each observation was left-out once, resulting in 19 total iterations. The best-fitting polynomial order was defined as one that produced the lowest median cross-validation error across all iterations.

### Model generalizability to basic visual tasks

We applied the same observer model to behavioral data from tasks mediated by acuity (6) and contrast sensitivity (26). The model was configured identically to what is described in the Model section above and the same model parameters were fit to behavioral data using BADS (94). To simulate the Neutral condition, attentional gain was not included in the model. Narrow SF and broad SF gain profiles were used to simulate all exogenous and endogenous attentional effects, respectively.

### Acuity

The modeling strategy for the acuity task is outlined in Figure S6. Landolt squares were inputted to the model with stimulus parameters that matched those described in (6). The squares were 1°-wide Landolt squares with a line thickness of 0.05°. Images were padded with 0.5° of empty space on each side to avoid edge artifacts. Model responses were computed for Landolt squares with a small gap (<1°) on the top or bottom. The Eucledian norm of the difference between responses indexed localization performance in the task. The model was evaluated at the eccentricity tested in the experiment (9.375°) and at 10 linearly spaced gap sizes (0-30 arcmin). We characterized the full psychometric function by interpolating between gap sizes. Interpolation was used to reduce computational load; similar psychometric functions were generated when the model was evaluated at finer intervals. The available signal for discrimination was scaled so that the maximum d′ equaled 2 and gap thresholds were quantified as the gap size needed to attain d′=1. For each attention type, 10 free parameters were fit to 14 gap thresholds (7 observers x 2 cueing conditions (Neutral, Valid)).

### Contrast sensitivity

The modeling strategy for the contrast sensitivity task is outlined in Figure S7. Tilted gratings (±45°) were inputted to the model with stimulus parameters that matched those described in (26). Gratings were windowed by a cosine function with a FWHM of 2°, had one of 6 SFs (0.5, 1, 2, 4 and 8 cpd) and were simulated at each of the four eccentricities tested (0°, 3°, 6° and 12°). We omitted the highest SF tested in (26) because it fell outside the range of SF subbands (0.5-8 cpd) used to simulate texture segmentation performance. Grating images were padded with 0.5° of empty space on each side to avoid edge artifacts.

To simulate the signal available to an observer in the orientation discrimination task, we computed the Eucledian norm of the difference between orthogonal gratings. This procedure was repeated for each grating SF and eccentricity. Model population responses were evaluated at 7 log-spaced levels of contrast that were interpolated to characterize the full contrast response function (Figure S7C). Similar contrast response functions were produced when the model was evaluated at finer contrast steps. We scaled the available signal by the magnitude of internal noise to yield stimulus discriminability (Decision mechanism). Because internal noise varies with SF (101), the available signal was scaled such that the maximum d′ at the fovea equaled 2 for each SF. Contrast thresholds were then determined as the level of contrast required to reach d′=1 and their inverse indexed contrast sensitivity. For each attention type, 10 free parameters were fit to 360 contrast thresholds (9 observers x 2 cueing conditions (neutral, valid) x 4 eccentricities x 5 SFs).

## Author Contributions

M.J., D.J.H. and M.C. conceived the project; M.J. implemented the model with input from D.J.H. and M.C.; M.C. provided all the data to be modelled; M.J. wrote the manuscript with guidance and supervision from M.C. All three authors edited the manuscript.

## Acknowledgements

This work was supported by National Institutes of Health RO1-EY019693 to M.C. We thank Michael Landy, Jonathan Winawer, Antoine Barbot, Hsin-Hung Li, as well as Antonio Fernández, Nina Hanning, Marc Himmelburg, Luke Huszar & Yong-Jun Lin and other members of the Carrasco lab for their helpful comments and discussion.

## Supplementary Information

**Figure S1.**
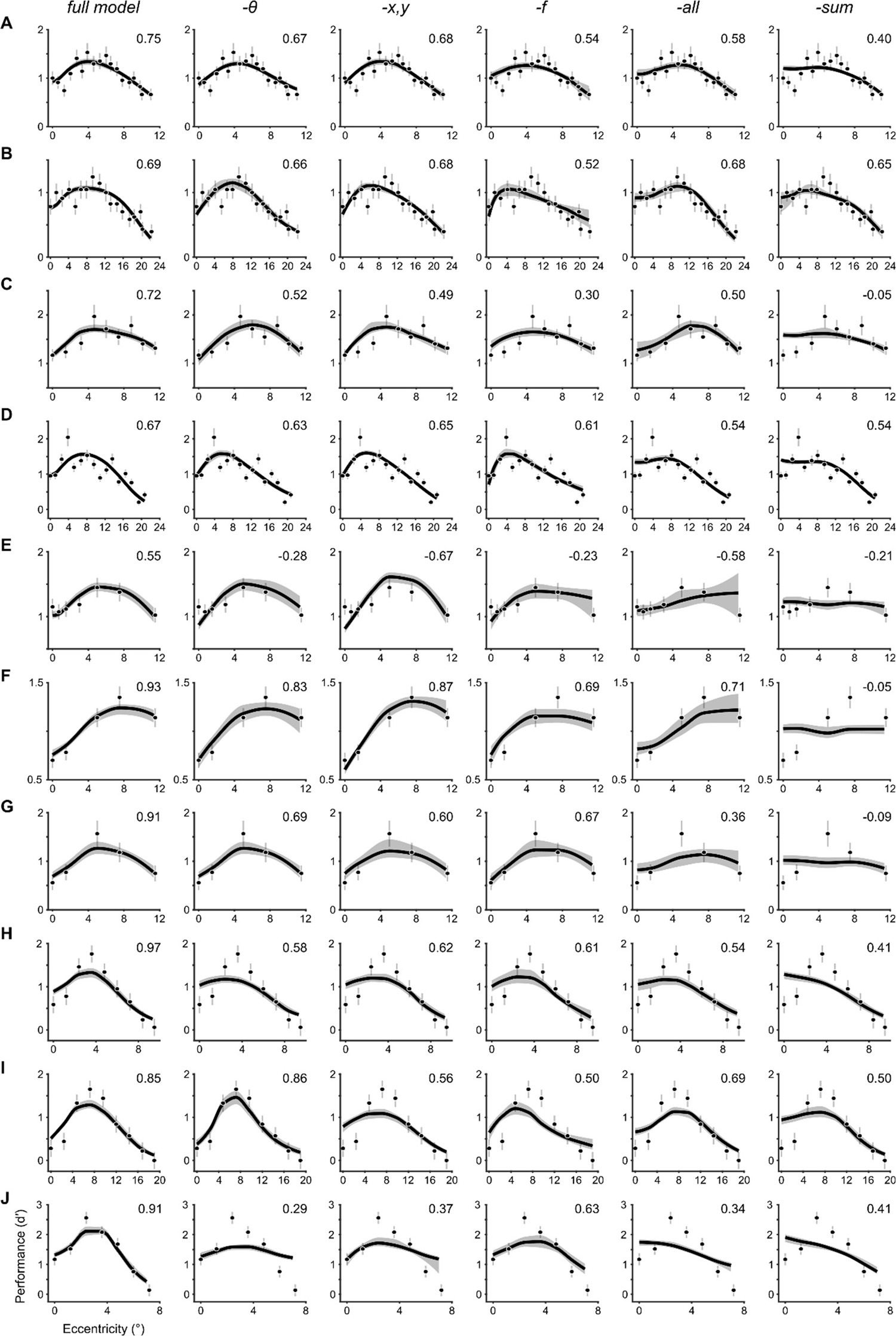
Model variants fit to the neutral condition of all texture segmentation experiments. Each row depicts the behavioral data from (A) Yeshurun & Carrasco, 1998 (27), Experiment 1; (B) Yeshurun & Carrasco, 1998 (27), Experiment 2; (C) Talgar & Carrasco, 2002 (29); (D) Carrasco, Loula & Ho, 2006 (30); (E) Yeshurun & Carrasco, 2008 (31); (F) Yeshurun, Montagna & Carrasco, 2008 (32), Experiment 2; (G) Yeshurun, Montagna & Carrasco, 2008 (32), Experiment 1; (H) Yeshurun, Montagna & Carrasco, 2008 (32), Experiment 3; (I) Yeshurun, Montagna & Carrasco, 2008 (32), Experiment 4; (J) Barbot & Carrasco, 2017 (33). Each column shows the fit of different model variants arranged in order of best-to-worst according to the model comparisons displayed in Figure 4D: ‘full’ denotes the full model, ‘-θ’ lacks cross-orientation suppression, ‘-x,y’ lacks surround suppression, ‘-f’ lacks cross-frequency suppression, ‘-all’ lacks all contextual modulation and ‘-sum’ lacks spatial summation. Dots and error bars denote group-average performance and ±1 SEM. The solid lines depict the median and shaded regions depict 68% confidence intervals of the bootstrapped distribution of model fits. Values in top-right of each panel denote the median R^2^ of the bootstrapped distribution of model fits. Negative R^2^ values indicate a model fit that captures less variance in the data than a horizontal line passing through the mean d′ across eccentricity.

**Figure S2.**
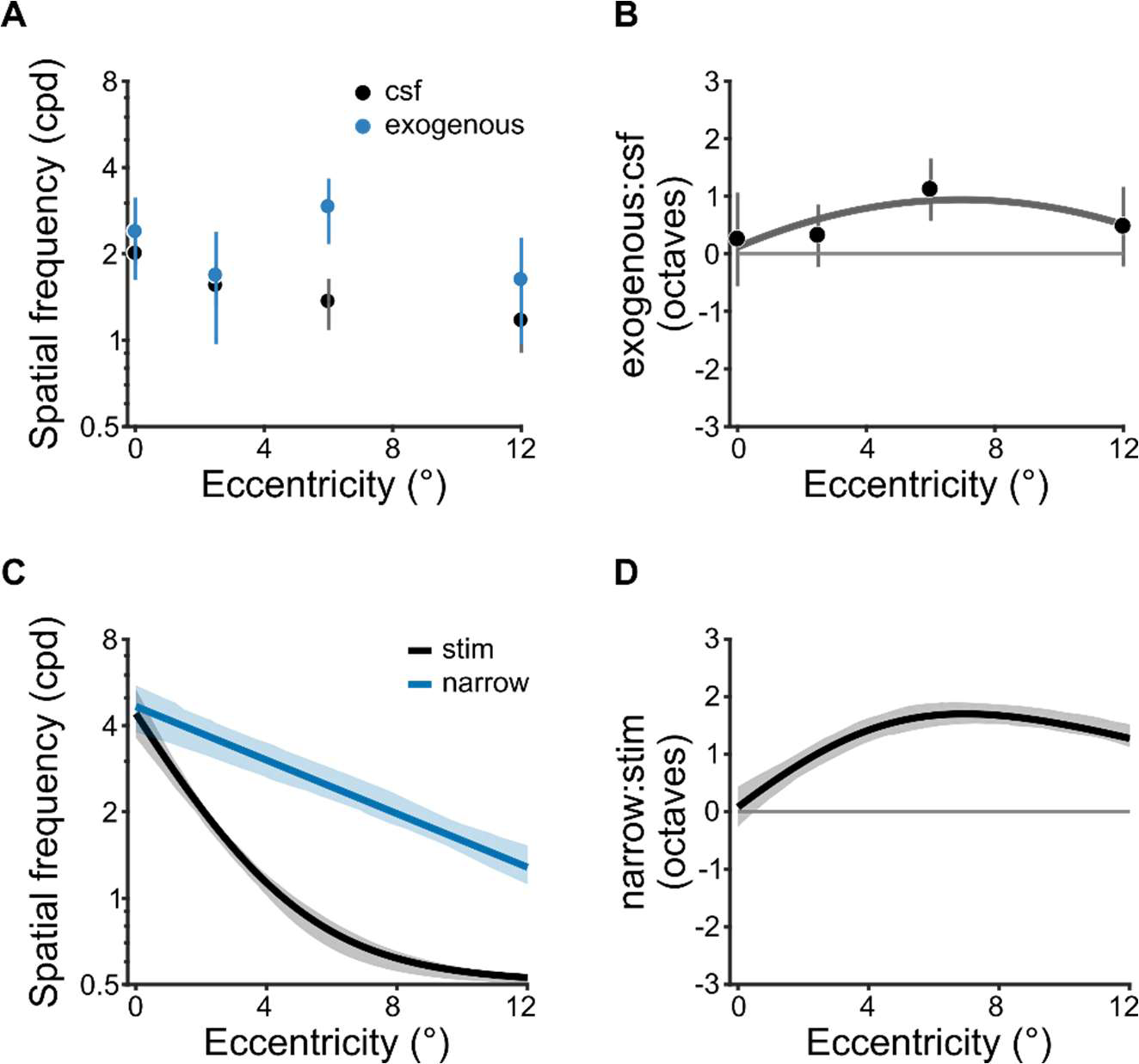
Spatial frequency operating range of exogenous attention. (A) Peak spatial frequency of baseline contrast sensitivity (CSF) and exogenous attentional modulation from Jigo & Carrasco, 2020 (26). Estimates were based on human contrast sensitivity, measured psychophysically with narrowband gratings. (B) Ratio (in octaves) of attentional and baseline peak spatial frequency tuning across eccentricity. Positive values denote an attentional preference for spatial frequencies higher than baseline. The solid line depicts the best-fitting second-order polynomial (i.e., parabola). Polynomial order was determined using leave-one-subject-out cross-validation (Methods, Cross-validation procedure). Dots in A and B depict group-average and error bars depict ±1 SEM. (C) Peak spatial frequency of the stimulus drive (stim) and the narrow SF attention gain profile (narrow). Estimates were derived from model fits to texture segmentation performance across all six exogenous attention experiments. (D) Ratio of the preferred spatial frequency for the stimulus drive and attentional gain. Solid lines indicate the median and shaded areas denotes 68% confidence interval of bootstrapped distribution in C and D.

**Figure S3.**
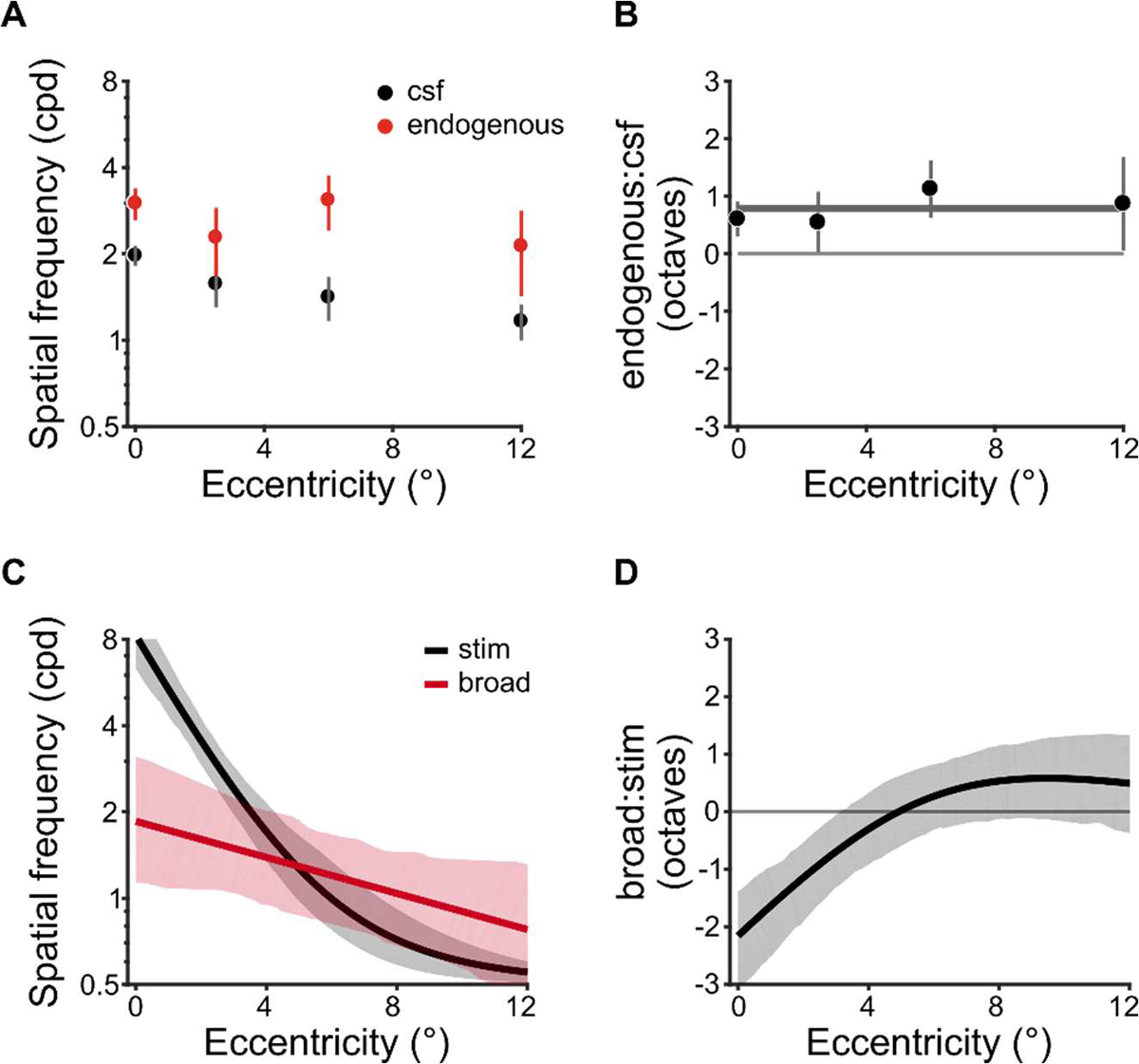
Spatial frequency operating range of endogenous attention. (A) The peak spatial frequency of baseline contrast sensitivity (CSF) and the center frequency of broad endogenous attentional modulation from Jigo & Carrasco, 2020 (26). Estimates were based on human contrast sensitivity, measured psychophysically with narrowband gratings. (B) Ratio (in octaves) of attentional and baseline and spatial frequency preferences across eccentricity. Negative values denote an attentional preference for spatial frequencies lower than baseline. The solid line depicts the best-fitting zero-order polynomial (i.e., constant). Polynomial order was determined using leave-one-subject-out cross-validation (Methods, Cross-validation procedure). Dots in A and B depict group-average and error bars depict ±1 SEM (C) The center spatial frequency of the stimulus drive (stim) and the broad attentional gain profile (broad). Estimates were derived from model fits to texture segmentation performance across all six endogenous attention experiments. (D) Ratio of the preferred spatial frequency for the stimulus drive and attentional gain. Solid lines indicate the median and shaded areas denote 68% confidence intervals of the bootstrapped distribution in C and D.

**Figure S4.**
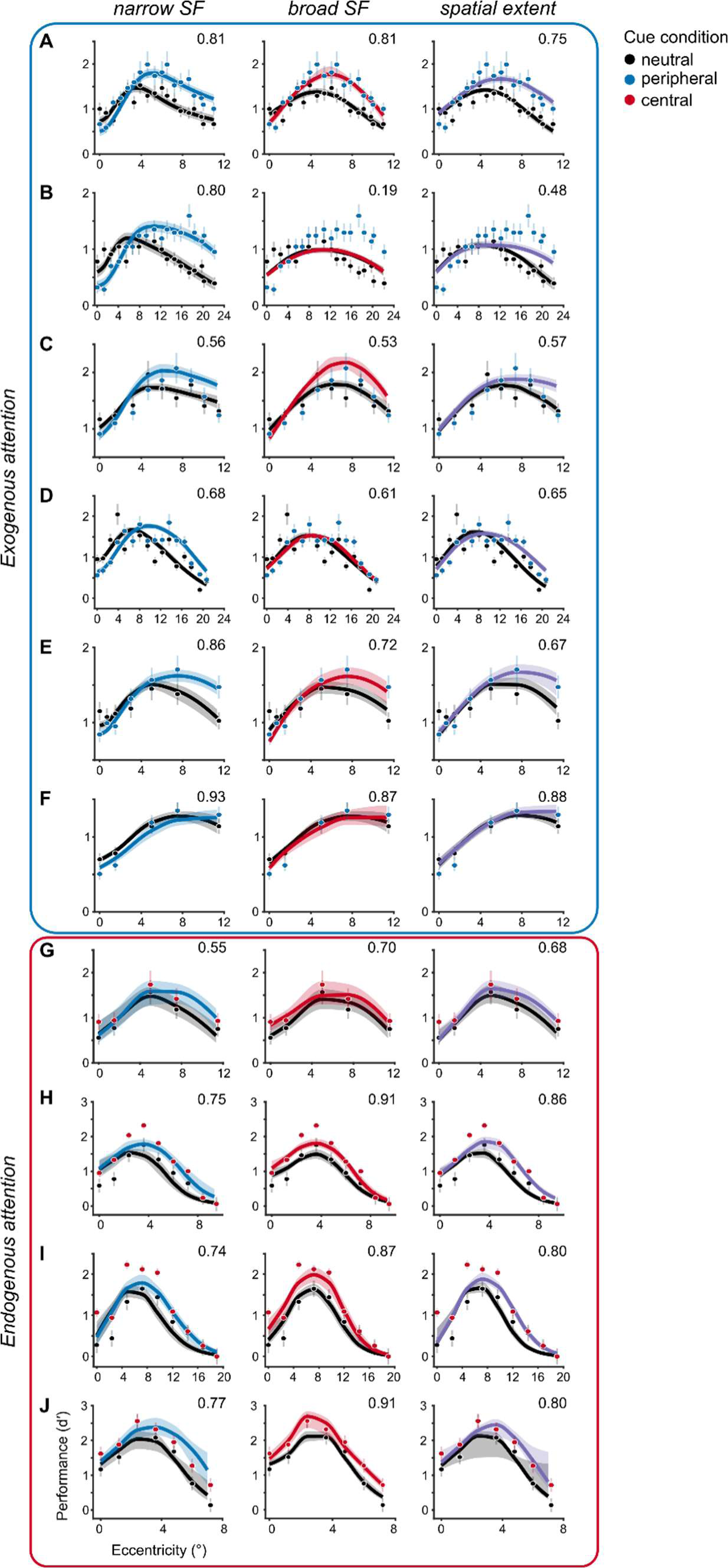
Attention model variants fit to behavioral data of all ten experiments. Each row depicts a different experiment: (A) Yeshurun & Carrasco, 1998 (27), Experiment 1; (B) Yeshurun & Carrasco, 1998 (27), Experiment 2; (C) Talgar & Carrasco, 2002 (29); (D) Carrasco, Loula & Ho, 2006 (30); (E) Yeshurun & Carrasco, 2008 (31); (F) Yeshurun, Montagna & Carrasco, 2008 (32), Experiment 2; (G) Yeshurun, Montagna & Carrasco, 2008 (32), Experiment 1; (H) Yeshurun, Montagna & Carrasco, 2008 (32), Experiment 3; (I) Yeshurun, Montagna & Carrasco, 2008 (32), Experiment 4; (J) Barbot & Carrasco, 2017 (33). Each column depicts a different attentional gain model. The numbers in the top-right of each panel denote the median R^2^ of the bootstrapped distribution of model fits.

**Figure S5.**
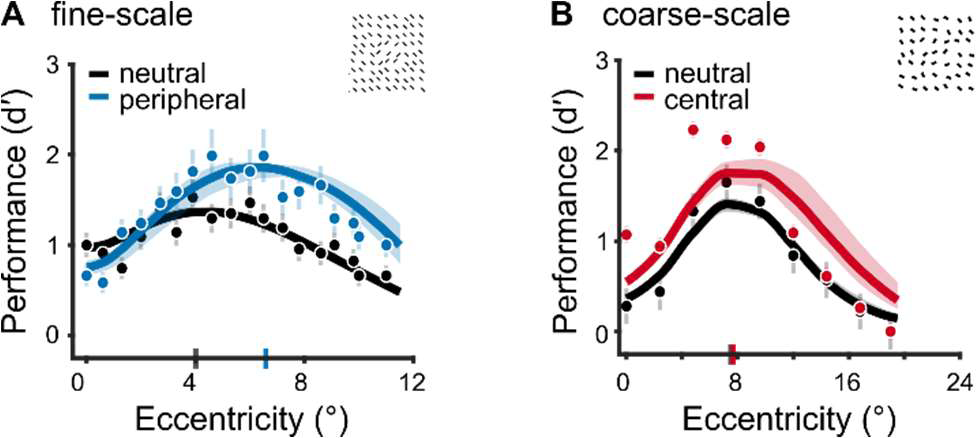
Model fits to jittered texture stimuli. (A) Predicted performance for Experiment 1 in Yeshurun & Carrasco, 1998 (27) using texture stimuli generated with line elements spatially jittered within the stimulus parameters of the experiment (Methods, Stimulus generation). An example jittered stimulus is shown in the top-right. Solid lines indicate the median and shaded regions depict 68% confidence intervals of the bootstrap distributions of model predictions. Gray and blue ticks on x-axis indicate peak of performance in the neutral and peripheral cueing condition, respectively. To generate the bootstrap distributions, model parameters were fixed to those that jointly captured all exogenous experiments. Then new jittered texture stimuli were input to the model on each iteration. (B) Predicted performance for Experiment 4 in Yeshurun, Montagna & Carrasco, 2008 (32) using texture stimuli generated with line elements whose orientation and spatial location were randomly jittered within the parameters of the experiment (Methods, Stimulus generation). Solid lines indicate the median and shaded regions depict 68% confidence intervals of bootstrap distributions of model predictions. Gray and red ticks on x-axis indicate peak of performance in the neutral and central cueing condition, respectively. To generate the bootstrap distribution, model parameters were fixed to those that jointly captured all endogenous experiments, then new jittered texture stimuli were input to the model on each iteration.

**Figure S6.**
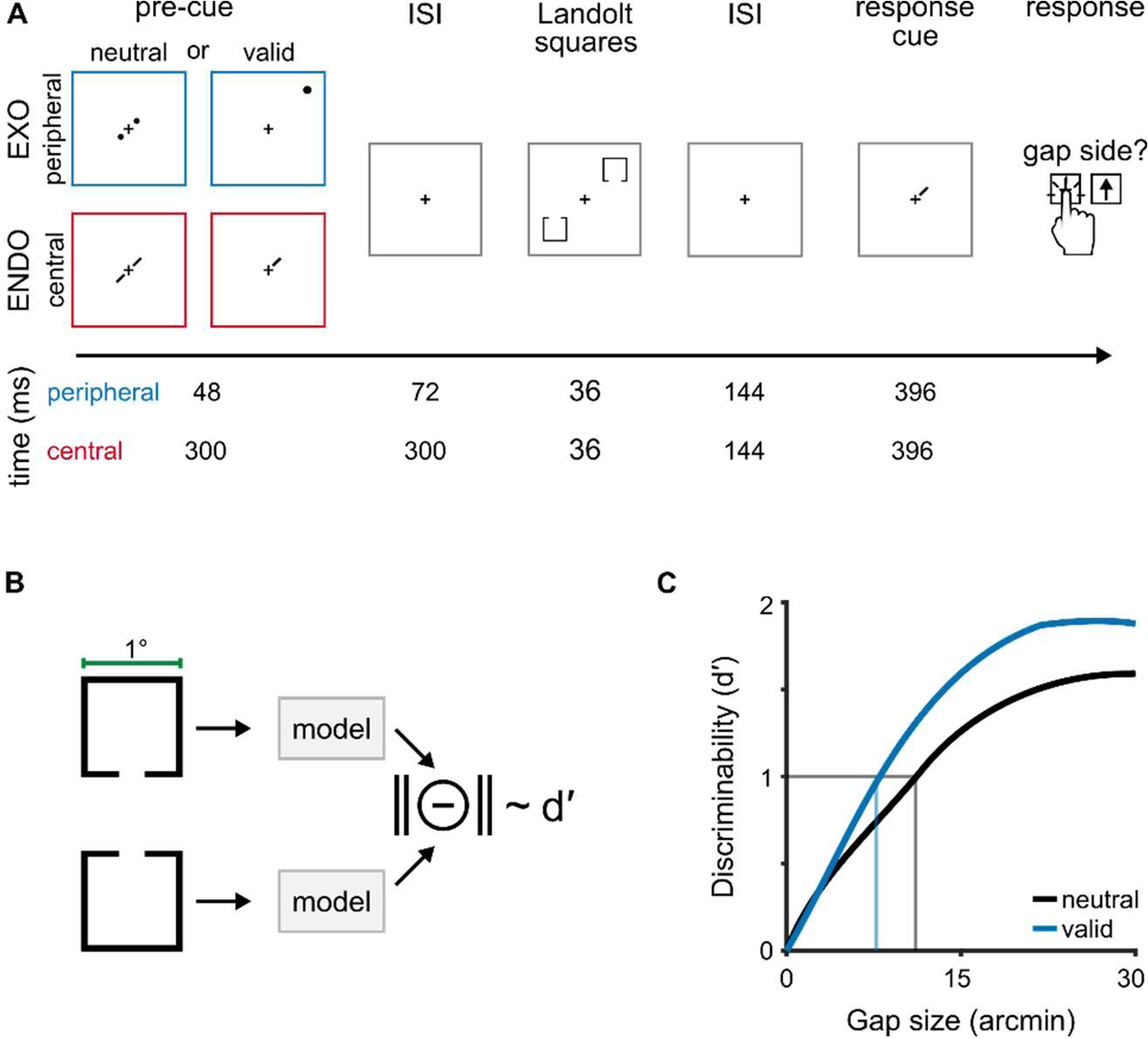
Behavioral protocol and modeling strategy for an acuity task. (A) Behavioral protocol adapted from (6). Observers performed a standard Landolt acuity task. The gap size in each 1°-wide Landolt square varied on a trial-by-trial basis and gap thresholds were measured in conditions where attention was distributed across both target locations (neutral) or focused at a single location (valid). Peripheral cues manipulated exogenous attention (EXO) whereas central, symbolic cues manipulated endogenous attention (ENDO). On each trial, two Landolt squares appeared on one of the two main diagonals of the visual field at 9.375°. Observers judged whether a gap appeared at the top or bottom of the Landolt square indicated by a response cue displayed at the end of the trial. The response cue equated uncertainty of the target’s location between neutral and valid cueing conditions. Gap thresholds indexed visual acuity in each condition. The timing information for peripheral (blue) and central (red) cueing conditions is given below each trial segment. (B) We modeled localization performance in this task by computing the discriminability (d′) between two Landolt squares, each with a gap at the top or bottom of the stimulus. (C) Model-derived discriminability across a range of gap sizes allowed for the creation of psychometric functions. To simulate neutral discriminability, attentional gain was not included in the model. We modeled discriminability for valid conditions with the narrow SF profile for exogenous attention and the broad SF profile for endogenous attention. Gap thresholds (vertical lines) were quantified as the gap size that resulted in d′=1 (horizontal line). We fit the model’s thresholds in each cueing condition to the behavioral data shown in Figure 9A-B.

**Figure S7.**
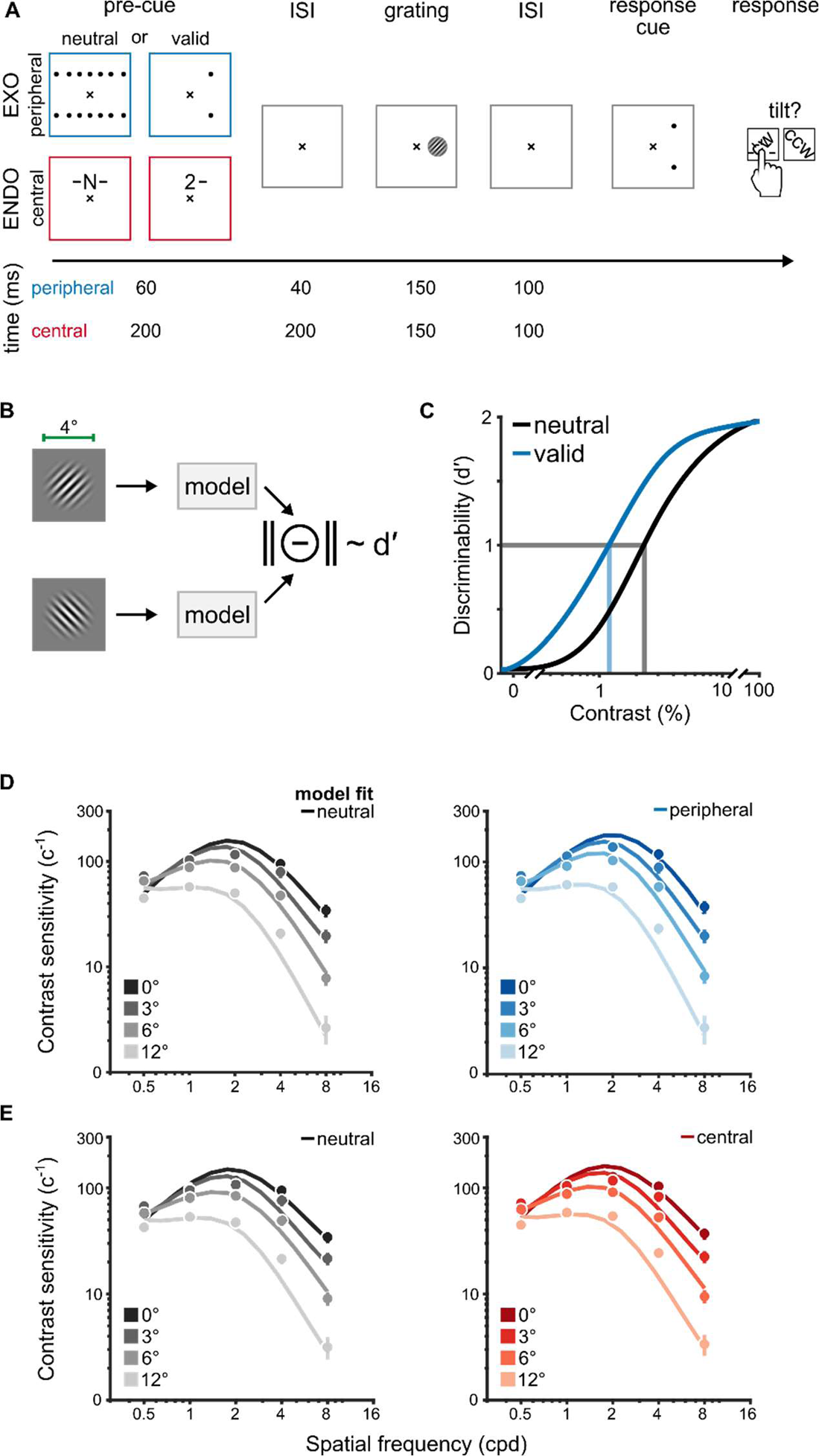
Behavioral protocol and modeling strategy for a contrast sensitivity task. (A) Behavioral protocol adapted from (26). Observers performed an orientation discrimination task on 4°-wide grating stimuli that varied in their contrast, SF and eccentricity. Grating contrast varied on a trial-by-trial basis and contrast thresholds were measured in conditions where attention was distributed across all possible target locations (neutral) or focused at a single location (valid). Peripheral cues manipulated exogenous attention (EXO) whereas central, symbolic cues manipulated endogenous attention (ENDO). On each trial, a single grating appeared along the horizontal meridian at 0°, 3°, 6° and 12° of eccentricity. The grating was tilted ±45° from vertical. After the onset of a response cue, observers judged whether the grating was oriented clockwise (CW) or counter-clockwise (CCW) from vertical. The response cue equated uncertainty of the target’s location between neutral and valid cueing conditions. The timing information for peripheral (blue) and central (red) cueing conditions is given below each trial segment. (B) We modeled orientation discrimination performance by computing the discriminability (d′) between two gratings, each tilted ±45° from vertical. Stimulus discriminability was simulated across a range of contrast levels for eccentricities and SFs tested in (26). (C) We simulated contrast response functions for each cueing condition. To simulate the neutral condition, attentional gain was not included in the model. Discriminability in the valid condition was modeled with the narrow SF profile for exogenous attention (i.e., peripheral cueing condition) and the broad SF profile for endogenous attention (i.e., central cueing condition). Contrast thresholds (vertical lines) were quantified as the level of contrast that resulted in d′=1 (horizontal line). The inverse of contrast thresholds indexed contrast sensitivity and were fit to the behavioral data. (D) Contrast sensitivity functions for neutral (left) and peripheral cueing conditions (right). The dots and error bars depict group-average contrast sensitivity and 68% confidence intervals for each eccentricity tested in (26). The solid lines are model fits to the behavioral data. (E) Contrast sensitivity functions for neutral (left) and central cueing conditions (right). Visualization conventions follow those in D. The vertical black lines in Figure 9C-D depict the peak SF of neutral contrast sensitivity functions, which indexed observers’ baseline tuning preferences. The ratio between valid and neutral contrast sensitivity indexed attentional effects across SF, shown in Figure 9C-D.

**Table S1.** Model parameters. Bolded entries indicate model components.

| Parameter | Description |
| --- | --- |
| **Stimulus drive** |  |
| $t_T$ | peak SF (log<sub>2</sub>-cpd) |
| $t_{min}$ | minimum preferred SF (cpd); *fixed at 0.5* |
| $m_T$ | SF tuning change across eccentricity (octaves/°) |
| $b_T$ | SF FWHM bandwidth (octaves) |
| **Contrast gain** |  |
| $g_\sigma$ | gain at fovea |
| $m_\sigma$ | slope along eccentricity |
| **Normalization pool** |  |
| $\delta_f$ | SF pool bandwidth (octaves); *fixed at 1* |
| $\delta_\theta$ | Orientation pool bandwidth (°); *fixed at 180* |
| $\delta_{pos}$ | Spatial pool width (°); *fixed at $\frac{2}{f}$* |
| **Spatial summation** |  |
| $\hat{\delta}_{pos}$ | Summation pool width (°); *fixed at $\frac{2}{f}$* |
| **Attentional gain profile** |  |
| $a_N$ or $a_B$ | attentional SF tuning at the fovea (log<sub>2</sub>-cpd) |
| $m_N$ or $m_B$ | slope of SF tuning across eccentricity (octaves/°) |
| $\gamma_N$ or $\gamma_B$ | amplitude |
| $b_N$ or $b_B$ | SF FWHM bandwidth (octaves) |
| $b_{pos}$ | spatial spread (°); *fixed at FWHM of 4* |

**Table S2.** Free parameters for the fits to the neutral condition of all ten texture segmentation experiments. The mapping between the experiment labels (a-j) and the respective references is given below the table and are consistent across all tables; italicized text describe the manipulation conducted in a given experiment. Bold values indicate the median and values within square brackets denote the 95% CI of the bootstrapped distribution of fitted parameters. min. = minimum; SF = spatial frequency; bw = bandwidth.

| parameter<br>description |  | Contrast gain |  | Stimulus drive |  |  |
| --- | --- | --- | --- | --- | --- | --- |
| | | $g_{\sigma}$ | $m_{\sigma}$ | $t_T$ | $m_T$ | $b_T$ |
|  |  | min. | slope | SF peak | SF slope | SF bw |
| Experiment | a | **2.3**<br>[2.2 2.5] | **-0.09**<br>[-0.09 -0.08] | **1.9**<br>[1.5 2.4] | **-0.8**<br>[-0.9 -0.6] | **1.7**<br>[1.6 2.0] |
|  | b | **2.3**<br>[2.1 2.5] |  |  |  |  |
|  | c | **2.7**<br>[2.5 2.8] |  |  |  |  |
|  | d | **2.7**<br>[2.6 2.8] |  |  |  |  |
|  | e | **2.2**<br>[2.0 2.5] |  |  |  |  |
|  | f | **2.7**<br>[2.4 2.8] |  |  |  |  |
|  | g | **1.9**<br>[1.5 2.3] |  | **2.8**<br>[2.4 3.1] |  |  |
|  | h | **1.8**<br>[1.7 2.0] |  |  |  |  |
|  | i | **2.6**<br>[2.5 2.8] |  |  |  |  |
|  | j | **2.7**<br>[2.6 2.8] |  |  |  |  |

<sup>a</sup> Yeshurun & Carrasco, 1998 (27). *Fine-scale texture; experiment 1.*
<sup>b</sup> Yeshurun & Carrasco, 1998 (27). *Coarse-scale texture; experiment 2.*
<sup>c</sup> Talgar & Carrasco, 2002 (29). *Target meridian: lower vertical.*
<sup>d</sup> Carrasco, Loula & Ho, 2006 (30). *Orientation discrimination: baseline adaptation.*
<sup>e</sup> Yeshurun & Carrasco, 2008 (31). *Attentional cue size: cue size 1.*
<sup>f</sup> Yeshurun, Montagna & Carrasco, 2008 (32). *Target meridian: horizontal; experiment 2.*
<sup>g</sup> Yeshurun, Montagna & Carrasco, 2008 (32). *Target meridian: horizontal; experiment 1*
<sup>h</sup> Yeshurun, Montagna & Carrasco, 2008 (32). *Fine-scale texture; experiment 3.*
<sup>i</sup> Yeshurun, Montagna & Carrasco, 2008 (32). *Coarse-scale texture; experiment 4.*
<sup>j</sup> Barbot & Carrasco, 2017 (33). *Target meridian: intercardinal; baseline adaptation.*

**Table S3.** Free parameters for the fits to neutral and peripheral cueing conditions of the six exogenous attention experiments. Bold values indicate the median and values in square brackets depict the 95% CI of the bootstrapped distribution of fitted parameters. min. = minimum; bw = bandwidth; amp. = amplitude.

| parameter<br>description |  | Contrast gain |  | Stimulus drive |  |  | Narrow gain profile |  |  |  |
| --- | --- | --- | --- | --- | --- | --- | --- | --- | --- | --- |
| | | $g_{\sigma}$ | $m_{\sigma}$ | $t_T$ | $m_T$ | $b_T$ | $a_N$ | $m_N$ | $b_N$ | $\gamma_N$ |
|  |  | *min.* | *slope* | *SF peak* | *SF slope* | *SF bw* | *SF peak* | *SF slope* | *SF bw* | *amp.* |
| Experiment | a | **1.9**<br>[1.7 2.2] | **-0.05**<br>[-0.07 -0.04] | **2.1**<br>[1.8 2.8] | **-0.7**<br>[-0.9 -0.5] | **1.6**<br>[1.4 2.1] | **2.3**<br>[1.8 3.0] | **-0.2**<br>[-0.3 -0.01] | **2.9**<br>[1.5 4.8] | **4.3**<br>[3.0 7.8] |
|  | b | **1.7**<br>[1.5 2.1] |  |  |  |  |  |  |  |  |
|  | c | **2.6**<br>[2.2 2.8] |  |  |  |  |  |  |  |  |
|  | d | **2.1**<br>[2.8 2.5] |  |  |  |  |  |  |  |  |
|  | e | **1.5**<br>[1.5 2.0] |  |  |  |  |  |  |  |  |
|  | f | **2.0**<br>[1.7 2.7] |  |  |  |  |  |  |  |  |

**Table S4.** Free parameters for the fits to the neutral and central cueing conditions of the four endogenous attention experiments. Bold values indicate the median and values within square brackets depict the 95% CI of the bootstrapped distribution of fitted parameters. min. = minimum; bw = bandwidth; amp. = amplitude.

|  |  | Contrast gain |  | Stimulus drive |  |  | Broad gain profile |  |  |  |
| --- | --- | --- | --- | --- | --- | --- | --- | --- | --- | --- |
| *parameter* | | $g_{\sigma}$ | $m_{\sigma}$ | $t_T$ | $m_T$ | $b_T$ | $a_B$ | $m_B$ | $b_B$ | $\gamma_B$ |
| *description* |  | *min.* | *slope* | *SF peak* | *SF slope* | *SF bw* | *SF peak* | *SF slope* | *SF bw* | *amp.* |
| **Experiment** | g | **1.6**<br>[1.5 2.1] | **-0.09**<br>[-0.1 -0.06] | **3.4**<br>[2.8 4.0] | **-0.8**<br>[-1.0 -0.5] | **1.3**<br>[1.1 1.6] | **1.7**<br>[0.5 2.3] | **-0.2**<br>[-0.3 -0.1] | **4.5**<br>[2.1 6.6] | **3.1**<br>[2.2 6.2] |
|  | h | **1.7**<br>[1.5 1.9] |  |  |  |  |  |  |  |  |
|  | i | **2.5**<br>[2.2 2.8] |  |  |  |  |  |  |  |  |
|  | j | **1.7**<br>[1.5 1.9] |  |  |  |  |  |  |  |  |

**Table S5.**
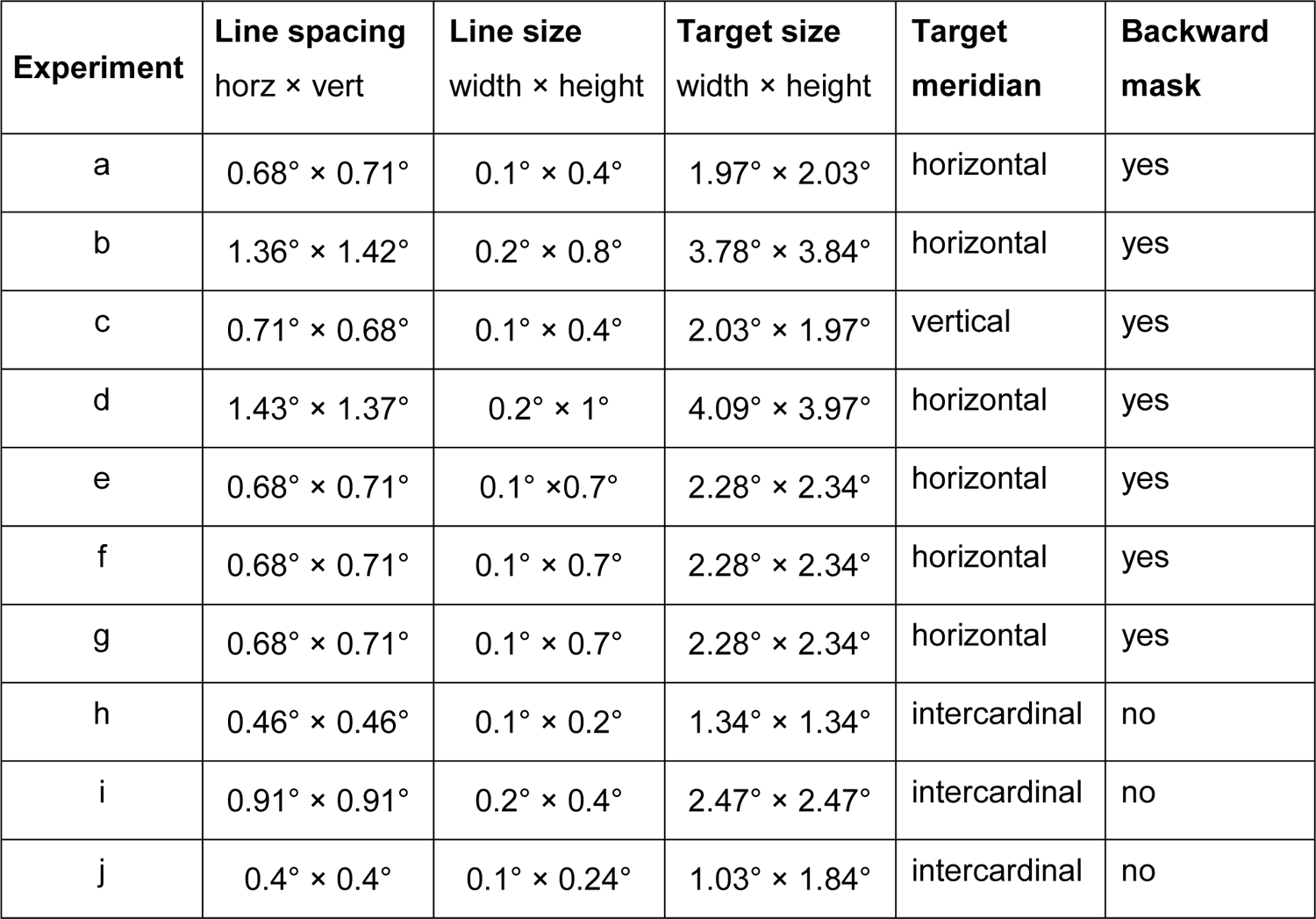
Stimulus parameters for each texture segmentation experiment.

| Experiment | Line spacing<br>horz × vert | Line size<br>width × height | Target size<br>width × height | Target<br>meridian | Backward<br>mask |
| --- | --- | --- | --- | --- | --- |
| a | 0.68° × 0.71° | 0.1° × 0.4° | 1.97° × 2.03° | horizontal | yes |
| b | 1.36° × 1.42° | 0.2° × 0.8° | 3.78° × 3.84° | horizontal | yes |
| c | 0.71° × 0.68° | 0.1° × 0.4° | 2.03° × 1.97° | vertical | yes |
| d | 1.43° × 1.37° | 0.2° × 1° | 4.09° × 3.97° | horizontal | yes |
| e | 0.68° × 0.71° | 0.1° × 0.7° | 2.28° × 2.34° | horizontal | yes |
| f | 0.68° × 0.71° | 0.1° × 0.7° | 2.28° × 2.34° | horizontal | yes |
| g | 0.68° × 0.71° | 0.1° × 0.7° | 2.28° × 2.34° | horizontal | yes |
| h | 0.46° × 0.46° | 0.1° × 0.2° | 1.34° × 1.34° | intercardinal | no |
| i | 0.91° × 0.91° | 0.2° × 0.4° | 2.47° × 2.47° | intercardinal | no |
| j | 0.4° × 0.4° | 0.1° × 0.24° | 1.03° × 1.84° | intercardinal | no |

**Table S6.** Best-fitting parameters for acuity and contrast sensitivity tasks. min. = minimum; bw = bandwidth; amp. = amplitude; acuity = acuity experiment (6); CS = contrast sensitivity experiment (26). The 95% confidence intervals show parameter values for fits to texture segmentation experiments, split by attention type.

| parameter<br>description |  | Contrast gain |  | Stimulus drive |  |  | Attentional gain |  |  |  |  |
| --- | --- | --- | --- | --- | --- | --- | --- | --- | --- | --- | --- |
| | | $g_{\sigma}$ | $m_{\sigma}$ | $t_T$ | $m_T$ | $b_T$ | a | m | b | $\gamma$ | $b_{pos}$ |
|  |  | min. | slope | SF peak | SF slope | SF bw | SF peak | SF slope | SF bw | amp. | spread |
| EXO | 95%<br>CI | [1.5 2.8] | [-0.07 -0.05] | [1.8 2.8] | [-0.9 -0.5] | [1.4 2.1] | [1.8 3.0] | [-0.3 -0.01] | [1.5 4.8] | [3.0 7.8] | - |
|  | Acuity | 2 | -0.07 | 1.8 | -0.8 | 1.4 | 2.8 | -0.03 | 1.5 | 8 | 0.6 |
|  | CS | 2.5 | -0.03 | 1.1 | -0.2 | 1.4 | 1.5 | -0.01 | 2.3 | 1.5 | 5.2 |
| ENDO | 95%<br>CI | [1.5 2.8] | [-0.1 -0.06] | [2.8 4.0] | [-1.0 -0.5] | [1.1 1.6] | [0.5 2.3] | [-0.3 -0.1] | [2.1 6.6] | [2.2 6.2] | - |
|  | Acuity | 1.5 | -0.09 | 3 | -0.9 | 1.1 | 2 | -0.3 | 5.3 | 15.0 | 0.6 |
|  | CS | 2.5 | -0.04 | 1.1 | -0.2 | 1.5 | 2.6 | -0.2 | 4.2 | 1.2 | 5.2 |

## Notes

### Competing Interest Statement

The authors have declared no competing interest.

### Summary of Updates

We add new simulations showing that model predictions generalize to attention effects on acuity and contrast sensitivity.

